# ChimericFragments: Computation, analysis, and visualization of global RNA networks

**DOI:** 10.1101/2023.12.21.572723

**Authors:** Malte Siemers, Anne Lippegaus, Kai Papenfort

**Author notes:** Corresponding author: Kai Papenfort, Friedrich Schiller University of Jena, Institute of Microbiology, General Microbiology, Winzerlaer Straße 2, 07745 Jena, Germany.

## Abstract

RNA-RNA interactions are key for post-transcriptional gene regulation in all domains of life. While ever more experimental protocols are being developed to study RNA-RNA interactions on a genome-wide scale, computational methods to analyze the underlying data are lagging behind. Here, we present ChimericFragments, an analysis and visualization framework for RNA-seq experiments producing chimeric RNA molecules. ChimericFragments implements a novel statistical method based on the complementarity of the base-pairing RNAs around their ligation site and is compatible with several widely used experimental procedures. We demonstrate that ChimericFragments enables the systematic identification of RNA regulators and RNA-RNA pairs and outperforms existing approaches.

## BACKGROUND

Base-pairing of two complementary RNA sequences is a fundamental principle of gene expression control in many, if not all, organisms. In eukaryotes, numerous different classes of non-coding RNAs have been described (*e.g.* microRNAs, long non-coding RNAs, and circular RNAs) controlling transcription, translation, or both [1, 2]. In bacteria, the majority of base-pairing regulators classify as small RNAs (sRNAs), which are ∼50-250 nucleotides in length and typically act together with RNA chaperones to recognize target transcripts through RNA duplex formation [3]. The consequences associated with successful base-pairing range from translation inhibition and transcript degradation to translation activation and increased protein synthesis [4, 5].

Comparative genomics and global transcriptome analysis have uncovered thousands of non-coding RNAs with usually unknown regulatory functions [6]. To close this gap, various experimental protocols have been developed capturing RNA-RNA interactions at a transcriptome-wide scale [7–10]. These tools typically rely on proximity-based ligation of two RNAs followed by high-throughput sequencing of the chimeric RNA molecules, allowing to infer regulatory interactions for annotated, as well as newly identified transcripts. For example, LIGR-seq (LIGation of interacting RNA followed by high-throughput sequencing) led to the discovery of small nucleolar (sno)RNAs interacting with mRNAs in human cells [11], whereas RIL-seq (RNA interaction by ligation and sequencing) revealed novel RNA-RNA interactions and sRNA regulators in bacteria [12, 13]. Other key technologies for global RNA interactome analysis are CLASH [14], SPLASH [15], and PARIS [16], all of which enable the genome-wide annotation of RNA-RNA pairs.

Although the above mentioned protocols differ in their experimental design, they all produce chimeric sequencing reads that can be analyzed through various bioinformatic tools. Previously, each method came with its own computational pipeline to detect and quantify RNA duplex formation, however, several new tools now provide a generic platform for data analysis. Specifically, RNA_NUE_ [17] and ChiRA [18] enhanced RNA duplex detection and data interpretation by computing the hybridization energy of putative RNA duplexes and by improving the mapping of short reads, respectively. However, these tools do not enable data visualization.

Here, we introduce ChimericFragments, a computational platform for the analysis and interpretation of RNA-RNA interaction datasets starting from raw sequencing files (Fig. 1). Our platform enables rapid computation of RNA-RNA pairs, RNA duplex prediction, and a graph-based, interactive visualization of the results. ChimericFragments employs a new algorithm based on the complementarity of chimeric fragments around the ligation site, which boosts the identification of *bona fide* RNA duplexes. When applied to a published dataset, ChimericFragments allowed the discovery a novel sRNA that controls virulence gene expression in the major human pathogen, *Vibrio cholerae*. ChimericFragments is implemented in *Julia* and available at: https://github.com/maltesie/ChimericFragments.

**Fig. 1.**
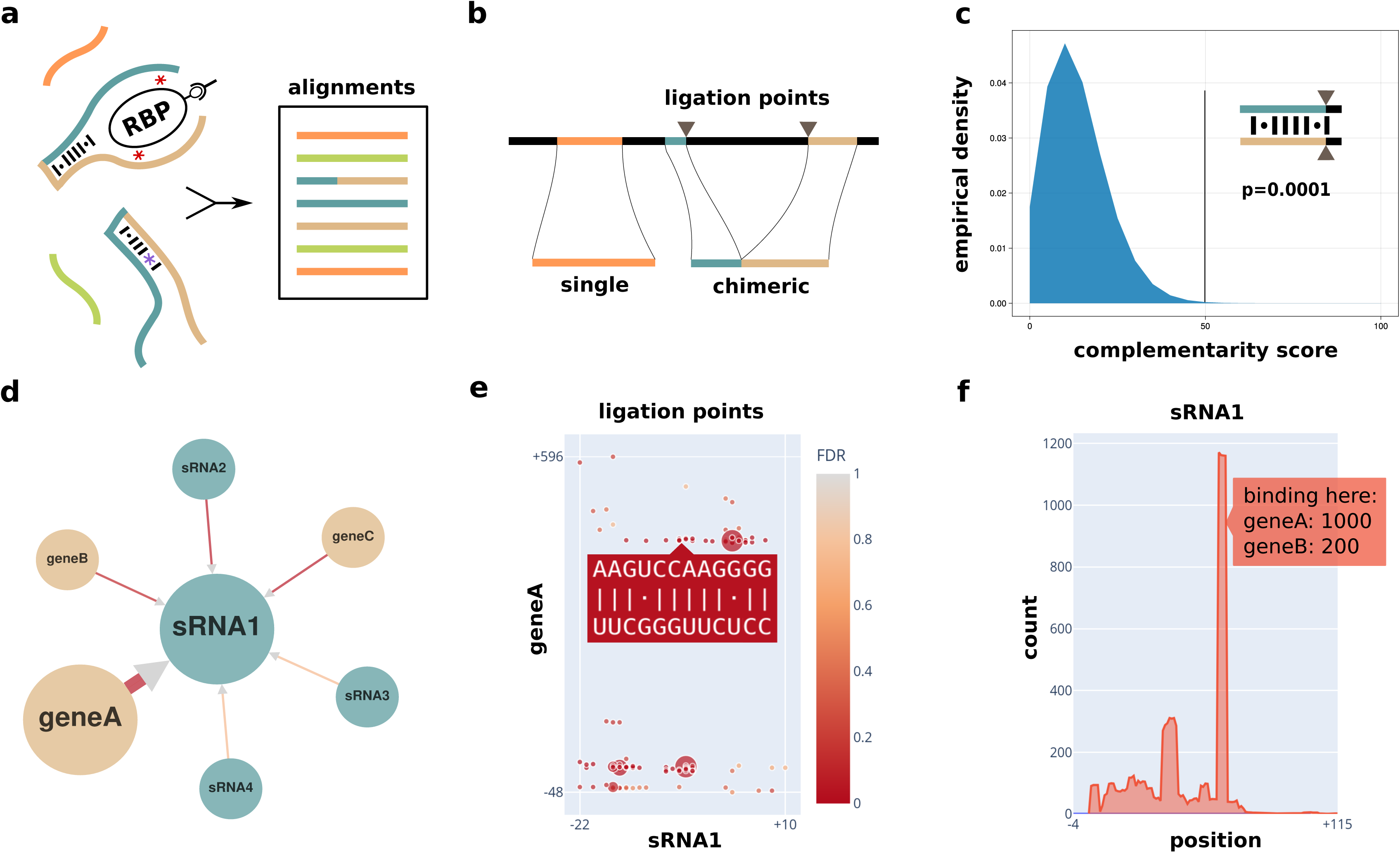
Graphical summary of the computational. (**a-c**) and the visual component (**d-f**) of ChimericFragments. **a**, Different experimental procedures (e.g. RIL-seq, CLASH, LIGR-seq) produce chimeric RNA molecules and sequence reads from them. Alignments of those reads are collected and sorted according to the position on the read they belong to. **b**, Alignments are classified to be single or chimeric and ligation points are saved. **c**, For each ligation point, the complementarity between the corresponding fragments is computed. A null model is computed for pairs of fragments from random positions on the genome and used to assign a *p*-value to ligated fragments. **d**, The global RNA-RNA network can be explored in an interactive graph-based visualization with color-coded annotation types and complementarity strength. **e**, Each ligation event can be inspected together with a basepairing prediction. **f**, All basepairing predictions for a selected transcript are visualized together and for each position, all partners are highlighted interactively.

## RESULTS

### ChimericFragments combines computational analysis and interactive visualization

Experimental workflows such as RIL-seq [13], CLASH [14], SPLASH [15], and PARIS [16] generate chimeric sequencing reads that are typically presented as large tables providing information on the identity, frequency, and statistical significance of the detected RNA-RNA interactions. However, this type of presentation fails to address the complex network structure associated with global RNA interactome studies and do not provide information on the positions of the relevant RNA duplexes [19]. ChimericFragments closes both of these gaps employing six main features (summarized in Fig. 1) and is compatible with a wide range of experiments producing chimeric RNA sequences.

The analysis of each dataset is split into three main categories: configuration, computation, and visualization. The configuration is set by defining the parameters in the template configuration file and all parameters are outlined in the supplied configuration template (default_config.jl). The computational part follows several main steps: 1^st^) preprocessing, 2^nd^) generation of a complete genome annotation, 3^rd^) split read alignment using bwa-mem2[20, 21], 4^th^) sorting, merging and classification of the produced alignments, and 5^th^) statistical evaluation of all interactions (Figs. 1a and b). For the latter, ChimericFragments captures ligation sites between fragments and computes the complementarity around them. This complementarity score is compared to a random model to produce p-values (Fig. 1c).

The resulting data are then used for a graph-based visualization displaying the annotated regions of the genome as nodes and the aggregate of all chimeras mapping to the same two nodes as edges (Fig. 1d). Additional interactive plots enable the prediction of RNA duplexes formation for every chimeric sequence displaying the frequency of the interaction, as well the position of RNA duplex formation relative to the annotation of the genes involved (Fig. 1e). Finally, the aggregate of all detected ligation sites is shown for each interacting transcript, allowing for the identification of preferred base-pairing sequences in regulatory RNAs and their targets (Fig. 1f). The visualization is implemented as a web application and can be used to share experimental results in the local network or over the internet. A detailed description of the control elements and the multiple data visualization modes in the graphical interface is provided in Figs. S1-4.

### Optimized mapping parameters increase the number of detected chimeras

A key step in the detection and analysis of base-pairing interactions from global RNA interactome studies is the mapping of chimeric sequencing reads to specific positions in the genome. Bwa-mem2 is an architecture-aware implementation of the bwa-mem algorithm, which can natively handle split alignments [20, 21]. We decided to use bwa-mem2 over other comparable algorithms, such as bowtie2 or STAR, based on former studies and due to its computational efficiency and precision [17, 22, 23].

To determine the suitability of bwa-mem2 for our approach, we set up several benchmarks for the aligner. Specifically, we quantified the sensitivity and precision of bwa-mem2 with respect to the size of the sequencing read and putative sequencing errors. Specifically, we generated several synthetic sequence libraries of different fragment lengths (15, 25 and 40 nucleotides) and with or without sequencing errors (Table 1). The libraries were each aligned with different values for two limiting parameters, *i.e.* seed length and minimum alignment score. The results show that without sequencing errors, bwa-mem2 aligns nearly all sequences of each length to the correct position (true positive rate, TPR ≥ 0.99, false positive rate, FPR ≤ 0.0005, Fig. 2a). A single sequencing error per read resulted in a dependency of the TPR and the FPR on the mapping parameters, as well as the length of the aligned sequence (Figs. S5a-f). We applied the same evaluation to another synthetic dataset of mixed length and an error probability of 0.5 per sequence (Figs. 2b-c). For this dataset, up to 82% of all chimeras were correctly aligned (highlighted in orange) with a FPR of 0.0119, while other combinations of seed length and minimum alignment score recovered 79%, 74%, and 67% of the chimeras with FPRs of 0.054, 0.0003, and 0.0002, respectively (shown in red, violet, and green). These findings highlight the importance of the alignment parameters for the detection of chimeric fragments, as well as a trade-off towards false discovery, which comes with less stringent parameters.

**Fig. 2.**
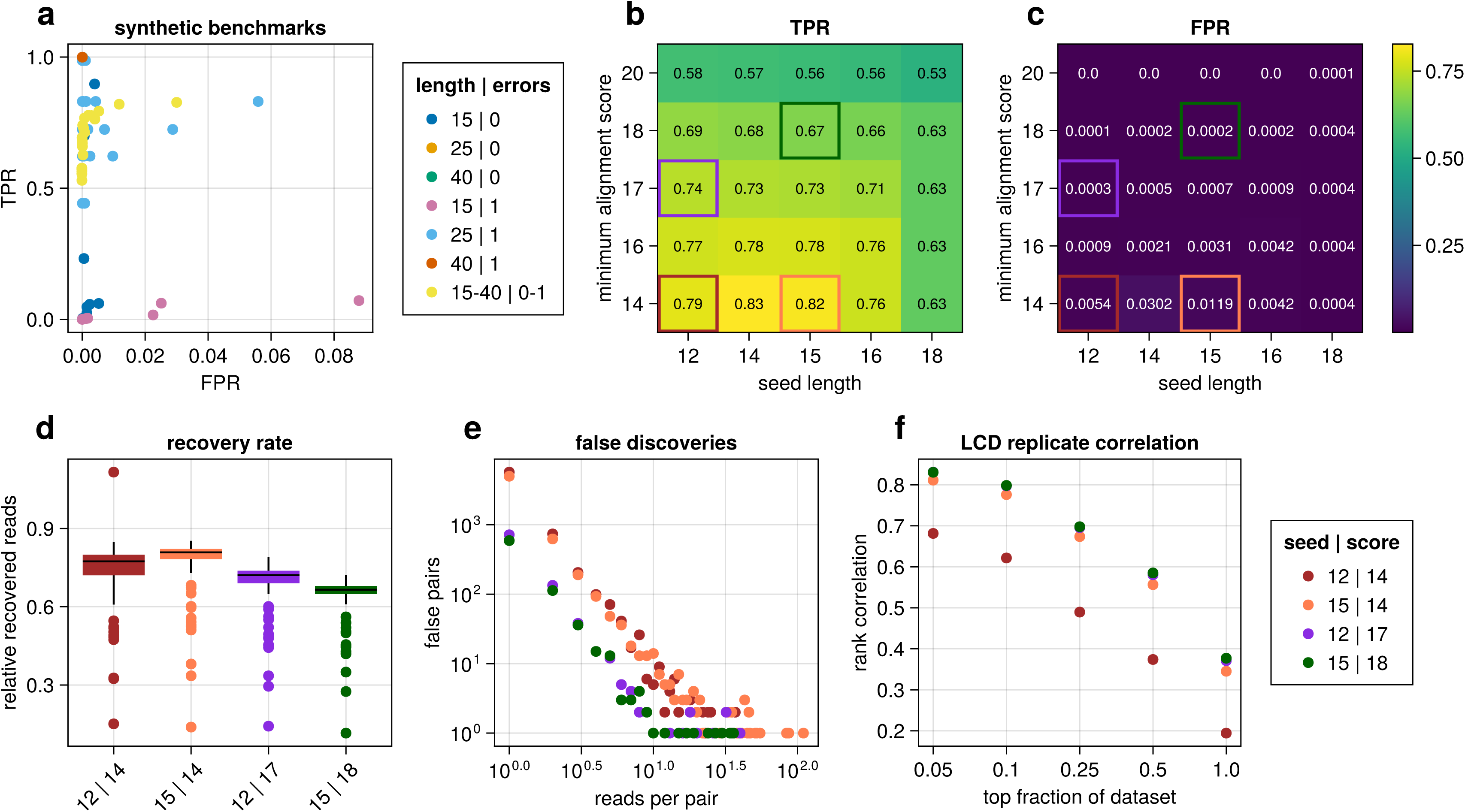
Benchmarking of bwa-mem2 for chimeric alignments. **a**, TPR and FPR of alignments in a synthetic dataset with combinations of *seed length* (12, 14, 15, 16, 18) and *minimum alignment score* (14, 16, 17, 18, 20). **b-c**, TPR and FPR for the mixed dataset (length 15-45, random mutation). Highlighted parameter combinations are used to compute the results shown in panels d-f. **d**, Box plots of recovery rates of reads per interaction in a synthetic chimeric dataset of 200 pairs of interacting regions with 1M randomly sampled reads. **e**, Falsely detected interactions in the evaluation of the synthetic chimera dataset (see d). **f**, Rank correlation of read counts per interaction for selected parameter combinations in 2 replicates of the RIL-seq dataset.

**Table 1.** Synthetic datasets to benchmark bwa-mem2 for chimeric alignments.

| fragment length | sequencing error |
| --- | --- |
| 15 | 0 |
| 15 | 1 |
| 30 | 0 |
| 30 | 1 |
| 45 | 0 |
| 45 | 1 |
| uniform random, 15-45 | uniform random, 0 or 1 |

We next tested the impact of the selected TPRs and FPRs on the analysis output. To this end, we generated a synthetic dataset of 200 pairs of interacting regions and sampled one million chimeras. We then analyzed this dataset using default parameters, only varying the seed length and the minimum alignment score. For the TPRs, the number of recovered chimeras per interacting pair almost perfectly matched the TPR found before (compare Figs. 2b and 2d). For the FPRs, we detected a high number of false chimeras, however, the vast majority came with drastically lower counts than the true chimeras (Fig. 2e). Few systematically false chimeras occurred in duplicated regions of the genome, which hindered bwa-mem2 to unequivocally map these reads.

To test our findings from the synthetic datasets with experimental data, we applied ChimericFragments to a previously published RIL-seq experiment containing two independent biological replicates [24]. Since no ground truth is available for these experiments, we used the correlation between the number of detected chimeric reads per interaction in the replicates as an indicator for the frequency of random misalignment. The Pearson correlation coefficient of the chimeric in the two replicates was consistently high (≥0.9, Fig. S5g). To get a more sensitive measure of our results, we also computed the rank correlation and found it to be very strong (≥0.9) for the top 10% fraction of the dataset, and decreasing when more interactions with lower read counts were added (Fig. 2f). All following analyses of experimental data were performed with a seed length of 12 and a minimum alignment score of 17, as this combination showed the best compromise between the TPR and the FPR in our synthetic benchmarks and was comparable to stricter parameters applied for the experimental data.

### Ligation points serve as indicators for stable RNA duplex formation

The above mentioned tools for global RNA interactome studies (*e.g.* RIL-seq, LIGR-seq, and CLASH) rely on the ligation of two proximal RNA molecules, resulting in the generation of chimeric sequencing reads [7, 25]. The general interpretation associated with the detection of a chimeric read is that the detected sequences base-paired, however, it remains unclear if these events describe spurious interactions, or stable RNA duplex formation. Previous work has addressed this problem by calculating the statistical significance of an interaction based on its frequency [17, 18, 26, 27] and initial attempts have been made to also consider additional parameters such as sequence complementarity and hybridization energy [17, 18], however, a statistical evaluation of those measures has not been performed.

To close this gap, ChimericFragments collects information on the two positions closest to the ligation site in a chimeric sequence, which we call the *ligation point* (Fig. 3a). For each chimeric read with a ligation point, the complementarity of the two fragments around the ligation point is computed. A complementarity score gets assigned to every chimera and its significance is evaluated by comparison to a model computed from the complementarity scores of randomly selected pairs of fixed sequences length from the genome. The distributions from the random model and the selected RIL-seq experiment overlap, however, clearly differ from each other (Fig. 3b). We next filtered our dataset based on the FDR assigned to each interaction and computed the correlation between the two biological replicates in the RIL-seq dataset. We observed a strong correlation (≥0.9) among the replicates for interactions with a FDR <0.01 and decreasing correlation for less stringent FDR-cutoffs (Fig. 3c), which resembled our previous analysis (Fig. 2f).

**Fig. 3.**
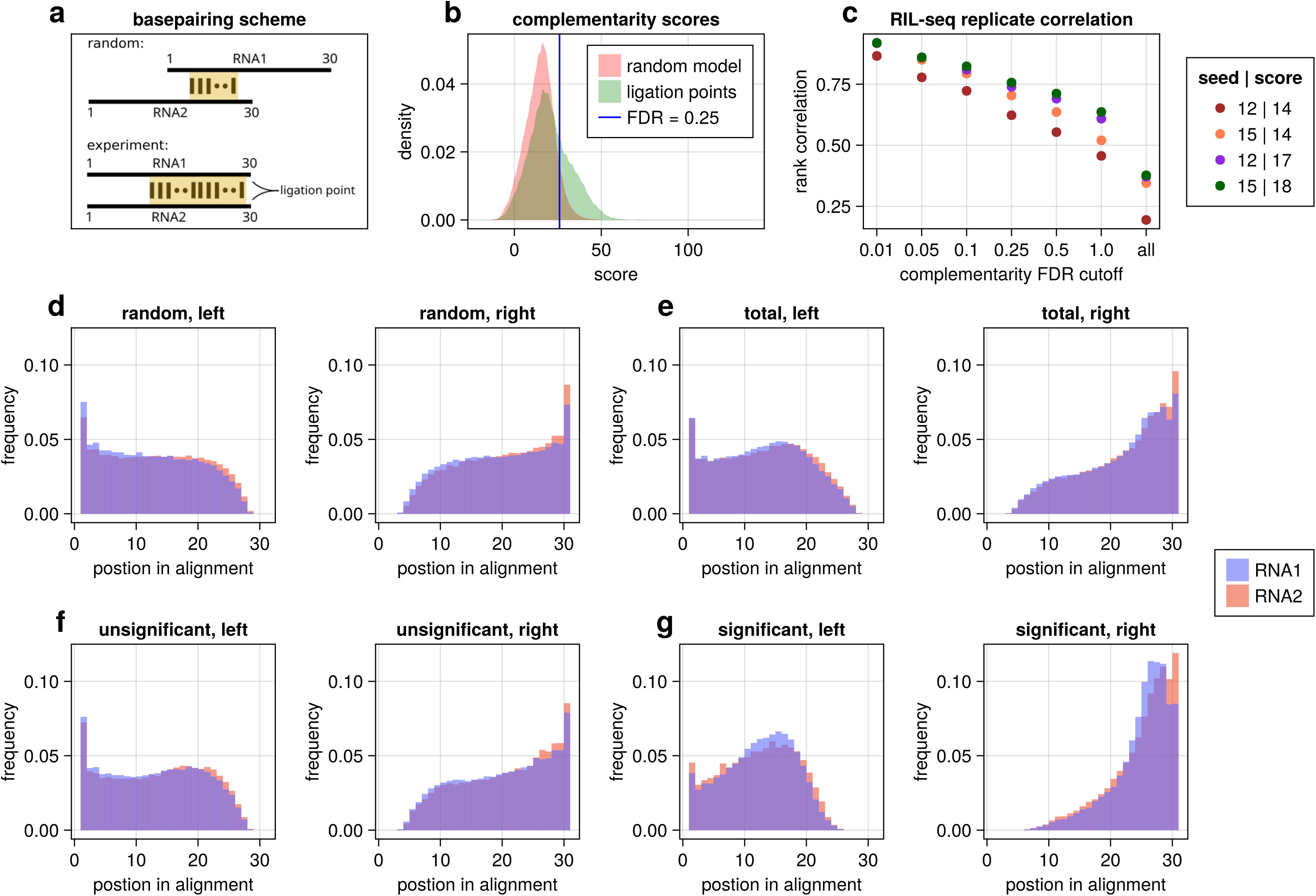
Statistics of basepairing predictions. **a**, Schematics of expected basepairing predictions between random (top) and experimentally derived sequence pairs **b**, Densities of score distributions for complementarity between random sequences (red) and sequences around ligation points (green) from a RIL-seq experiment, with a fixed length of 30 nucleotides. **c**, Rank correlation of read counts per interaction for selected mapping parameter combinations in 2 replicates of a RIL-seq experiment, filtered with respect to the significance of basepairing predictions **d**, Alignment ends histogram for alignments of random sequences of length 30. **e**, Alignment ends histogram for all alignments in a RIL-seq experiment. **f**, Alignment ends histogram for unsignificant (FDR *≤* 0.25) alignments in panel e. **g**, Alignment ends histogram for significant (FDR *>* 0.25) alignments in panel e.

To further understand the effect of filtering interactions based on their complementarity score, we analyzed the distribution of the complementary regions in their respective sequences. In the random model, the ends of the interacting sequences are distributed symmetrically (Fig. 3d). Using the significance of the complementarity score as a separation criterion, the total distribution of predicted RNA duplexes around ligation points splits into two populations (Fig. 3e), with the non-significant part closely resembling the distribution in the random model (Fig. 3f). The distribution of the significant interactions shows a strong preference for RNA duplex formation close to the ligation point (position 30, Fig. 3g, right) and the left ends of complementarity regions follow a left-tailed distribution with a peak around position 15 (Fig. 3g, left). In contrast, comparison of the significance values from our statistical evaluation and the commonly used Fisher exact test showed no association between the two methods (Figs. S6a-b). These data indicate that the ligation point of a chimeric sequence is a powerful indicator for the identification of stable RNA duplexes in global RNA interactome studies.

To benchmark our approach, we applied ChimericFragments to seven published datasets, involving six different organisms and three different experimental pipelines, *i.e. V. cholerae* (RIL-seq [24]), *Escherichia coli* (*E. coli*, RIL-seq and CLASH [13, 28]), enteropathogenic *E. coli* (RIL-seq [29]), *Pseudomonas aeruginosa* (RIL-seq [30]), *Salmonella enterica* (RIL-seq [31]), and *Bacillus subtilis* (LIGR-seq [32]). In all cases, ChimericFragments recovered significantly more interactions than initially reported in the respective studies (Figs. S6c-i). Of note, although our results are difficult to compare to the previously reported interactions due differences in the statistical methods to filter the datasets, we discovered ligation points for the majority of interactions in all studies. Except for the LIGR-seq dataset, ChimericFragments revealed more interactions with significant complementarity around the ligation points, when compared to the respective initial study.

We also analyzed two previously published RIL-seq datasets [13, 24] using the ChimericFragments pipeline. These datasets were collected in two different model organisms (*E. coli* and *V. cholerae*), allowing us to compare our results with 93 published and experimentally verified RNA-RNA interactions (56 for *E. coli* and 37 for *V. cholerae* [24, 33]). We detected relevant chimeras for 55 and 35 of these interactions, respectively (Table S1). In *E. coli*, ChimericFragments successfully predicted the reported interaction in 40 cases (∼73%) and similar numbers were obtained in *V. cholerae* (22/35, ∼63%). For comparison, RNAnue detected 19/55 (∼35%) interactions in *E. coli* and 15/35 (∼43%) interactions in *V. cholerae* (Table S1). Of note, ChimericFragments also comes with an improved runtime requiring 10 s/Mio. reads, whereas RNA_NUE_ required 1,290 s/Mio. reads (Fig. S7a).

### ChimericFragments reveals hidden sRNA-target mRNA pairs

To evaluate the ability of ChimericFragments to predict undetected RNA duplexes in global RNA interactome studies, we reanalyzed published RIL-seq datasets to search for unknown interaction with high complementarity scores. Specifically, we allowed read counts as low as 3 to apply our complementarity-based test of significance and investigated the complementarity in low frequency interactions. When compared to previous work [24] (using a minimal cutoff of 20 reads per interaction and a Fisher exact test FDR<0.05), our optimized mapping parameters increased the number of RNA pairs by ∼12% (3,580). Dropping the requirement of significance according to the Fisher exact test, the number of interactions further improved by ∼2.8-fold (8,976), whereas chimeras with read counts of ≥3 and significant base-pairing predictions increased by ∼4.1-fold (12,917) (Fig. 4a). In summary, we detected a total of 58,261 interactions (>3 reads) of which 32,890 (∼56.5%) contained a ligation point, resulting in 22,897 (∼39.3%) significant RNA duplexes (FDR ≤0.25).

**Fig. 4.**
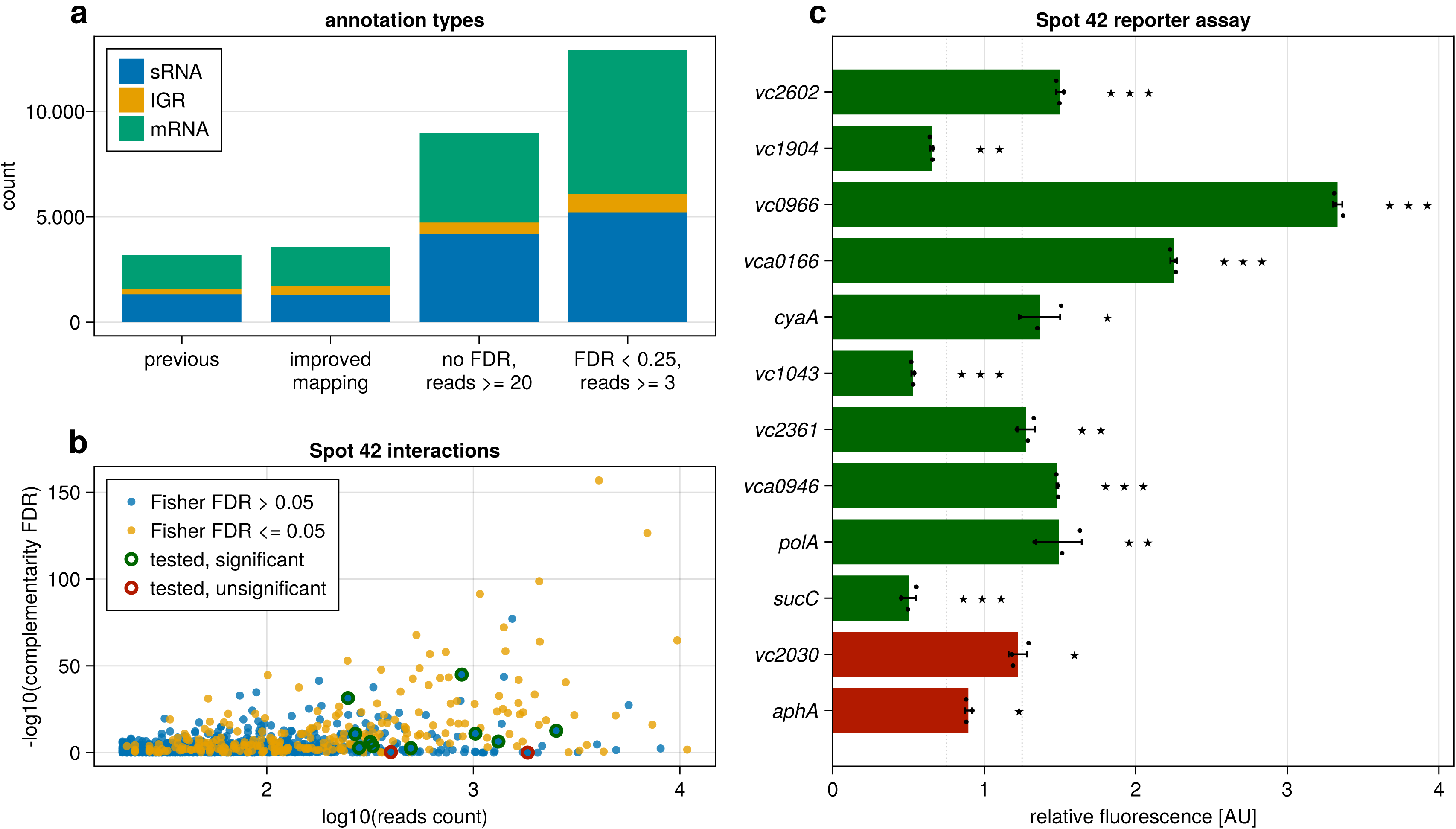
Improved characterization of known regulators. **a**, Comparison of amounts of interactions recovered in the RIL-seq dataset with our previously published analysis and ChimericFragments. **b**, Significance of basepairing predictions for all interactions with more than 20 reads. Colors indicate interactions detected with our previous analysis (violet) and additional interactions detected only with ChimericFragments (orange). Red circles highlight interactions picked for validation. **c**, Translational GFP reporter fusions were cotransformed with a constitutive Spot 42 expression plasmid or an empty control plasmid in *E. coli* Top10 cells and GFP production was measured. Green bars reflect significant (FDR ≤ 0.25) and red bars unsignificant (FDR > 0.25) basepairing. Bars show the mean of the measurements in three independent biological replicates. For each target, all measurements were divided by the mean of the control measurements and error bars are equal to the respective propagated uncertainty. Significance (unpaired t-test) of the difference towards the control samples is indicated by stars: *: p *≤* 0.05, **: p *≤* 0.01, ***: p *≤* 0.001

To corroborate these findings, we selected 10 putative new targets of the well-studied Spot 42 sRNA [34], showing FDR values >0.05 in the Fisher exact test (Fig. 4b), and tested their regulation using an *in vivo* post-transcriptional reporter assay [35]. Eight of these targets also showed significant RNA duplex formation and for all these targets we confirmed a regulatory effect >25% (Fig. 4c). In contrast, the two remaining target did not display regulation by Spot 42.

### ChimericFragments provides insights into the mechanisms of sRNA-mediated gene regulation

Bacterial sRNAs frequently employ multiple base-pairing sequences to interact with target mRNAs, which adds to their function as global regulators of gene expression[4, 36]. However, the identification of base-pairing sequence elements in a given sRNA is typically not straight-forward based on conservation analysis alone [37–39]. ChimericFragments addresses this problem as it computes RNA duplexes for every chimeric RNA pair with a ligation point, which are visualized in a summary plot (Fig. 1f). Single peaks indicate one base-pairing sequence in the sRNA (Fig. S7b), or the target (Fig. S7c), whereas multiple peaks predict more than one base-pairing sequence (Fig. 5a). Using this strategy, we discovered 20 sRNAs with a single base-pairing sites in *V. cholerae* and 35 that contained two or more sites (Figs. S7e-f).

**Fig. 5.**
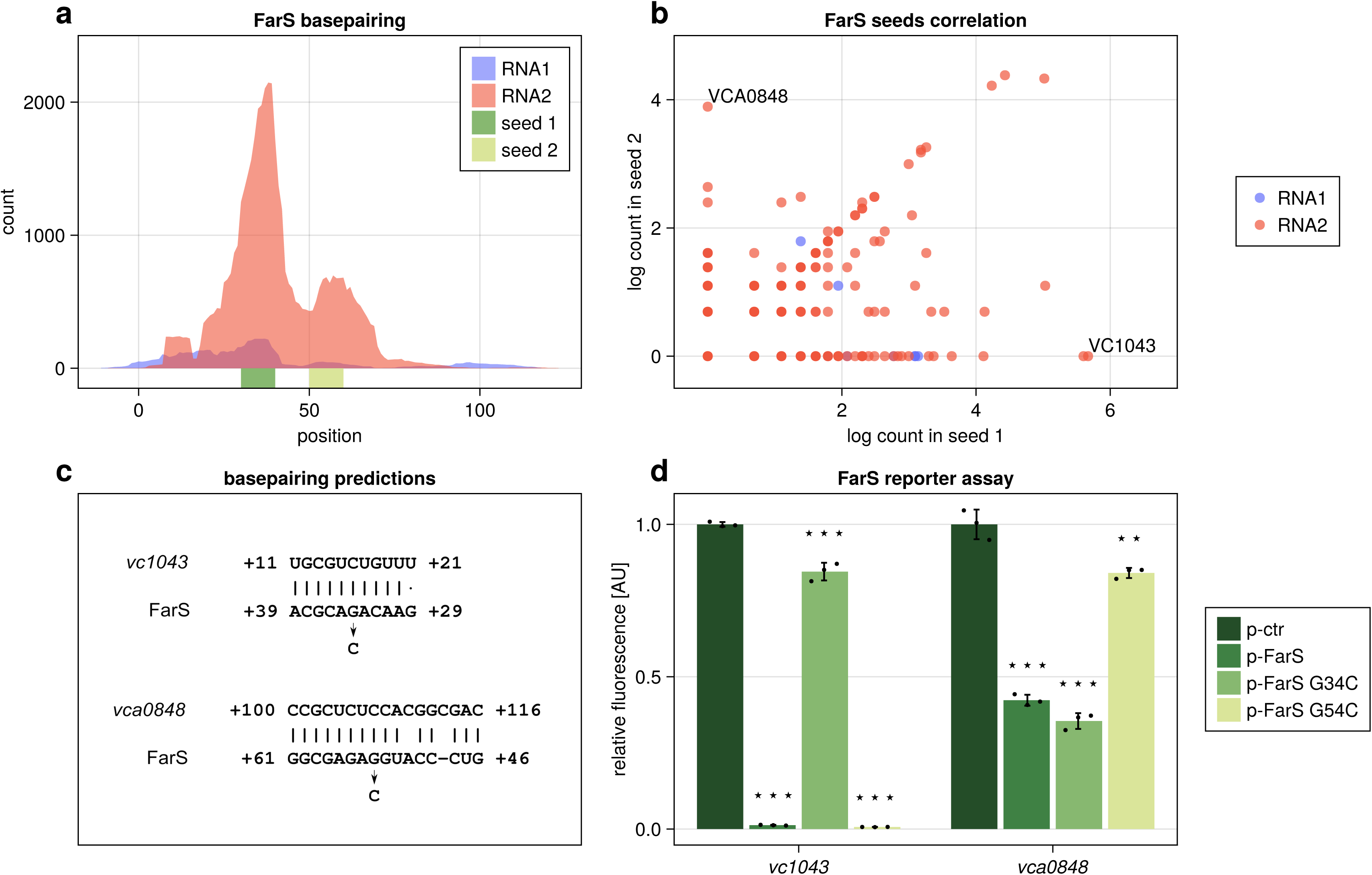
Improved characterization of known regulators. **a**, Two seed regions in FarS (seed 1, dark green; seed 2, light green) detected by aggregation of all basepairing predictions of all its targets (FDR *≤* 0.25). **b**, Log transformed counts of reads supporting complementarity in seed1 or seed2 in FarS for all predicted targets (FDR *≤* 0.25, 1 pseudocount added). **c**, Basepairing prediction between FarS seed 1 and *vc1043* (top) and FarS seed 2 and *vca0848* with bases for nucleotide exchange marked. **d**, Translational GFP reporter fusions for *vc1043* and *vca0848* together with an empty control plasmid (ctr) or FarS expression plasmids (FarS, G34C and G54C) were cotransformed in *E. coli* Top10 and GFP production was measured. Bars show the mean of the measurements in three independent biological replicates and error bars are equal to the respective standard deviation. For both targets, all measurements were divided by the mean of the control measurements. Mutations G34C and G54C correspond to the seed regions marked with the respective color in panel a. Significance (unpaired t- test) of the difference towards the control samples is indicated by stars: **: p *≤* 0.01, ***: p *≤* 0.001

The summary plots also classify the results depending on the position of a fragment in a sequencing read relative to its partner (RNA1 precedes RNA2; Fig. 3a). These data can inform a potential mode of regulation as targets preferentially occupy the first position, whereas regulators (*e.g.* sRNAs) are frequently found in the second position [13, 40]. Indeed, when analyzed by our ChimericFragments pipeline, we confirmed that Hfq-binding sRNAs were more frequently recovered as RNA2, when compared to their target mRNAs (Fig. S7g).

To test if this information would allow us to discover new sRNA targets, we focused on the FarS sRNA, which was previously shown to use a single base-pairing site to control two related fatty acid degradation genes [41]. In contrast, our data suggested FarS regulates additional genes using two base-pairing sites (Fig. 5a). To validate these predictions, we focused on two target mRNAs with high frequency exclusive to each region: *vc1043* (encoding a fatty acid transporter [42]), which interacts with a novel base-pairing sequence in FarS and *vca0848* (encoding a GGDEF family protein [43]), employing the previously reported base-pairing site (Figs. 5b-c). Post-transcriptional reporter assays revealed that *vc1043* and *vca0848* are both repressed by FarS, however, introduction of single nucleotide mutations (G34C and G54C; Fig. 5c) confirmed that base-pairing is specific to the predicted base-pairing site (Fig. 5d). Taken together, our analysis show that ChimericFragments generates testable hypothesis that enable a better understanding of the molecular mechanisms underlying post-transcriptional gene regulation.

### Identification and characterization of novel regulatory RNAs using ChimericFragments

Regulatory RNAs and target mRNAs have distinct properties in global RNA networks. Whereas regulatory RNAs often interact with hundreds of targets, mRNAs mostly interact with one or few sRNAs, but not with other mRNAs [3, 4, 12, 44]. We used this difference in the local network structure to search for undiscovered sRNAs in intergenic regions (IGRs) of the genome. To this end, we computed the network of all interactions between coding sequences (CDSs) and IGRs, revealing two IGRs pairing with numerous putative target mRNAs (Fig. S8). We further analyzed the IGR with the most targets (located between the *vc0715* and *vc0719* genes; Fig. 6a) and found that most targets shared a predicted binding site within the *vc0715*::*vc0719* IGR (Fig. 6b). To support our hypothesis for regulatory function of the *vc0715*::*vc0719* IGR, we inspected published transcriptome datasets for a potential sRNA transcript [45, 46]. Indeed, these analyses revealed a ∼100 nt long transcript, which we named NetX (network derived RNA), and we validated its expression by Northern blot analysis (Fig. 6c).

**Fig. 6.**
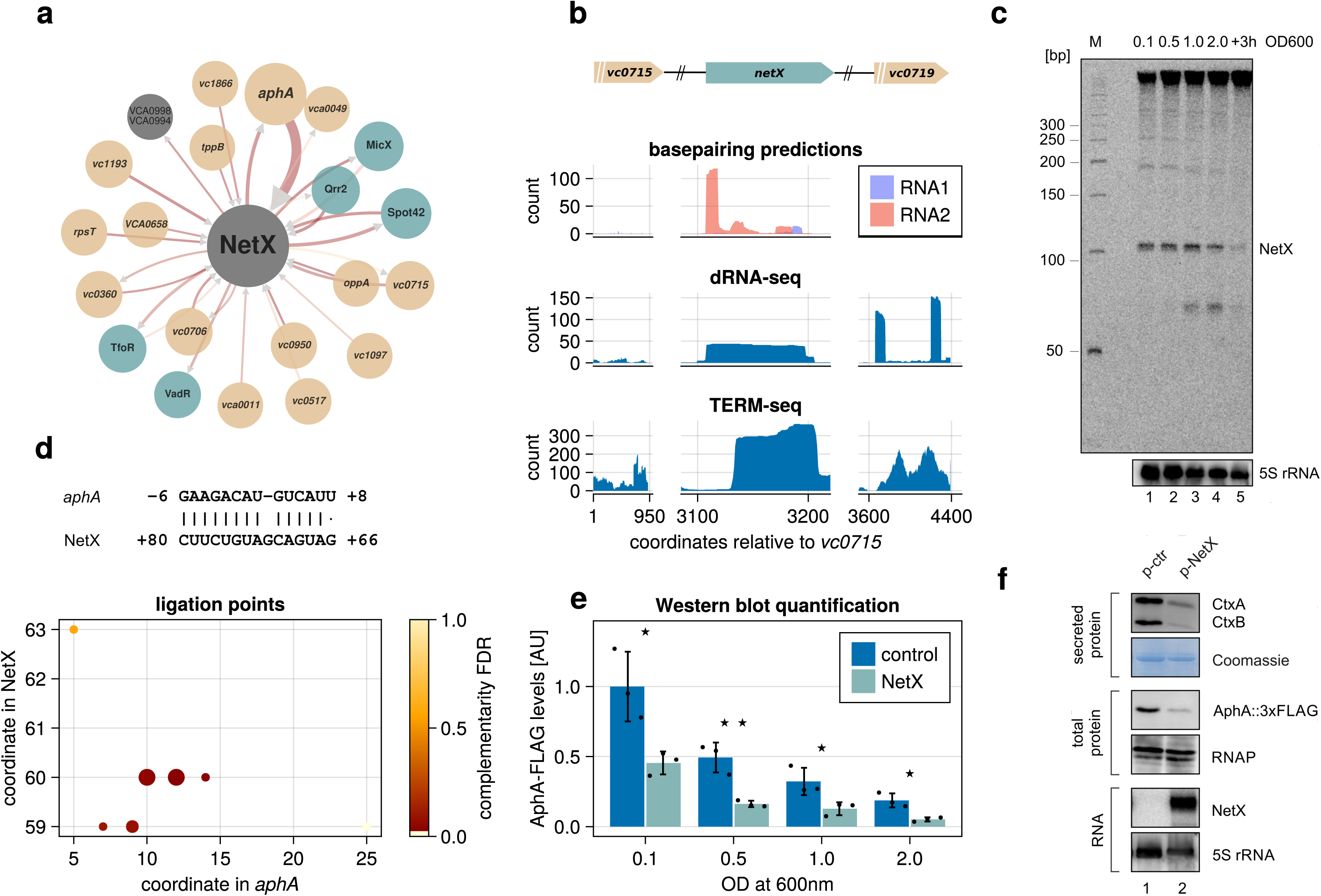
Identification and characterization of the NetX small RNA. **a**, Graph of all interactions of NetX with more than 3 reads captured by RIL-seq. **b**, Genomic context of the newly characterized sRNA NetX with aggregation of basepairing predictions for NetX and flanking genes together with coverage from a dRNA-seq experiment and a TERM-seq experiment at the same location. **c**, Northern blot detecting a transcript made at the region shown in panel b. RNA samples from *V. cholerae* wild-type cells were collected at various stages of growth. 5S ribosomal RNA served as a loading control. **d**, ChimericFragments plot of ligation points (bottom) between NetX and the interaction partner *aphA* with a selected basepairing prediction shared by all ligation points in the lower left corner of the plot (top). **e**, Quantification of Western blots comparing protein levels of AphA between WT and overexpression of NetX. Protein samples from *V. cholerae* wild-type cells carrying a chromosomal 3XFLAG in the *aphA* gene were collected at various stages of growth. Western Blot analysis was performed to measure AphA levels. Bars show the mean of three independent biological replicates and error bars are equal to the respective standard deviation. Significance (unpaired t-test) of the difference towards the control samples is indicated by stars: *: p *≤* 0.05, **: p *≤* 0.01 **f** *V. cholerae* wild-type cells carrying a chromosomal 3XFLAG-tag in the *aphA* gene were cultivated in AKI medium. Secreted protein, total protein and RNA samples were collected, and RNA and protein samples were monitored respectively by Northern and Western blot analysis. Coomassie staining, RNAP and 5S ribosomal RNA served as a loading control for Western and Northern blots, respectively.

We next focused on the regulatory role of NetX. According to ChimericFragments the most abundant target mRNA of NetX is *aphA*, encoding a key regulator of quorum sensing, biofilm formation, competence, and virulence in *V. cholerae* [47–50]. Our analysis predicted strong base-pairing between NetX and *aphA* (Fig. 6d) and Western blot analysis showed that over-expression of NetX resulted in reduced AphA protein levels in all stages of growth (Fig. 6e). Given the documented role of AphA in virulence gene expression and pathogenesis of *V. cholerae* [51], we extended our analysis and monitored the effect of NetX expression on cholera toxin (CtxAB) production. In line with our previous data, NetX strongly reduced CtxAB levels (Fig. 6f). Taken together, our results show that ChimericFragments allows the detection of novel RNA regulators and supports the hypothesis-driven research into their regulatory and biological roles in the cell.

## DISCUSSION

Base-pairing between two RNA molecules often depends on RNA chaperones such as Hfq and ProQ in bacteria, or SM-like proteins from eukaryotic and archaeal organisms [52–54]. Mutation of their respective genes typically impairs RNA duplex formation and in the case of Hfq it has been associated with pleiotropic phenotypic alterations, including defects in virulence gene expression in pathogenic bacteria [55]. Therefore, studying the molecular processes underlying RNA chaperone activity and global RNA-RNA interactions patterns is not only an important aspect of fundamental research, but also has implications for medicine and public health [56].

The past few years have brought a revolution in our understanding of how RNA-RNA interactions form at a global level due to the development of various new sequencing-based technologies [7, 9, 10]. In contrast to previous approaches, which frequently relied on the identification of individual RNA duplexes and/or the characterization of single RNA regulators, these technologies have paved the way to simultaneously analyze the interactomes of dozens to hundreds of regulatory RNAs and thousands of RNA-RNA pairs [7]. However, computational pipelines addressing the complexity of these datasets are scarce and it is often unclear how the detected interactions translate into functionally important discoveries [6].

ChimericFragments offers a computational framework that can help to overcome these limitations (Fig. 1). Specifically, the integrated graphical interface allows visualization of global RNA-interactomes, which can provide important information on the relevance of individual regulators or RNA-RNA interactions in the network. Previous work has shown that cellular RNAs constantly compete for interaction with RNA binding proteins, shaping the biophysical and biochemical parameters underlying post-transcriptional gene regulation [57, 58]. Thus, to understand the regulatory principles underlying RNA network performance, it is crucial to determine the structure of the network and key regulatory players involved [59]. Of note, ChimericFragments also enables the comparison of two or more network states allowing to study the regulatory features driving RNA network dynamics.

The evolution of regulatory RNAs is highly dynamic and, when compared to their protein counterparts, only poorly understood [36, 60–62]. Regulatory RNAs can be expressed from IGRs, as well as from the 5’- and 3’UTRs of mRNAs, and the coding sequence [63]. In addition, RNA regulators can also originate from stable transcripts, such as tRNAs [64, 65]. Therefore, the identification of base-pairing regulators from the pool of all cellular transcripts can be difficult based on standard transcriptome data. ChimericFragments allows the discovery of base-pairing regulators based on the number and quality of the interactions (Fig. 6). We demonstrate this feature of ChimericFragments through the identification and characterization of NetX, which we show is a previously unknown regulator of virulence gene expression in *V. cholerae*.

Our approach also revealed a second new sRNA regulator, named NetY, which is expressed from the IGR between the *vcr069* and *vc1803* genes (Fig. S9a). NetY accumulates as a ∼80 nucleotide long sRNA (Fig. S9b) and alike NetX base-pairs with various transcripts (Fig. S9c). However, in contrast to NetX (Fig. 6a), the majority of NetY’s interaction partners are other non-coding RNAs, suggesting that this sRNA might act as an RNA sponge. RNA sponges base-pairs with and inhibit the activity of non-coding regulators and are ubiquitous in prokaryotic and eukaryotic systems [66, 67]. Further investigations towards the mechanism underlying NetY-mediated base-pairing supported its role as a sponge RNA as the vast majority of chimeras contained NetY on the first position of the sequencing read (indicated in blue; Fig. S9c), which is a hallmark of target mRNAs and sponge RNAs [13, 26, 40]. In contrast, analogous analyses focusing on FarS and NetX revealed that their corresponding transcripts are typically found in the second position of the sequencing reads (indicated in red; Figs. 5a and 6b), suggesting their primary function is to regulate other transcripts. In the case of FarS, we also discovered that the sRNA contains two base-pairing sequences to interact with target transcripts and we also discovered multiple base-pairing sequences in various other sRNAs (Fig. S7a).

These analyses also identified RNA regulators that carry signatures of both categories, *i.e.* they seem to act as sponges when base-pairing with one set of targets, while in other interactions they likely function as the regulator. Of note, the base-pairing sequence involved in these interactions can be either overlapping (*e.g.* see VSsrna24; Fig. S9d) or occupy separate segments of the sRNA (*e.g.* see GcvB; Fig. S9e). The latter case could indicate a switch in the regulatory function of an sRNA depending on the use of a specific base-pairing sequence, which has not been previously observed. Again, these results highlight the strength of ChimericFragments in generating data-driven hypotheses that can be tested experimentally.

Finally, we designed ChimericFragments to be compatible with various experimental setups that have been used to detect RNA-RNA in bacteria (*e.g.* RIL-seq, CLASH, and LIGR-seq). However, ChimericFragments can also deal with data from eukaryotic organisms, as we demonstrate for a CLASH experiment from *Saccharomyces cerevisiae* (Fig. S9f; [68]). However, we note that larger genome sequences together with the relatively small size of eukaryotic microRNAs, siRNAs, and piRNAs will reduce the number of uniquely mapping sequencing reads in our pipeline, which complicates downstream analysis. Therefore, adjustments in the mapping strategy would be required to adjust ChimericFragments to these alternative datasets.

## CONCLUSIONS

In this study, we present ChimericFragments, a computational pipeline enabling the analysis and visualization of global RNA interactome data starting from raw sequencing data. ChimericFragments comes with several new features, including rapid detection of RNA-RNA interactions, prediction of the associated RNA duplexes, and global network visualization that together will facilitate the analysis and interpretation of such complex datasets. ChimericFragments also allows for the identification of previously overlooked RNA regulators, which in the case of NetX we demonstrate to have important roles in gene regulation and physiology. Our approach is compatible with several major experimental pipelines and thus has the potential to inform new biology beyond the realm of a single organisms or regulator.

## METHODS

### Genome annotation

ChimericFragments tags each alignment with an annotation. To capture all chimeras in a dataset, it relies on a fully annotated genome for both forward and reverse strands. The quality of the results depends on the quality of the annotation, so it is recommended to supply ChimericFragments with an accurate annotation of noncoding RNAs and 5’ and 3’ UTRs. The automatic annotation of the genome requires a set of CDS for which the regions up- and downstream are extended up to a maximum length or until another annotation is reached. The remaining regions of the genome will then be annotated for each strand as IGRs named by their flanking genes.

### Reads preprocessing

As a first step in the analysis, ChimericFragments uses fastp to preprocess the raw sequencing reads [69]. fastp removes adapters from the reads and read ends with low quality get trimmed and short reads discarded.

### Chimeric alignments with bwa-mem2

ChimericFragments uses bwa-mem2 [21] to map reads to the genome. To benchmark bwa-mem2 in its ability to detect chimeric alignments, several synthetic datasets were generated. All of them comprise of chimeric sequences of different length and either with a mutation at a random position or without. The datasets are summarized in Table 1.

### Annotation of alignments

Every aligned fragment is annotated uniquely. To efficiently find the annotation with the largest overlap, a binary interval tree is constructed to find m overlaps in a set of n annotations in *O*(*m* + log *n*). The annotation overlapping the most with the alignment is used to tag it with a name and a type and in cases of tied overlaps, the annotation with the lowest left coordinate on the genome is chosen.

### Sorting, merging and classification of alignments

All alignments are sorted first with respect to the read they come from and second to the order of their origin on the read sequence from 5’ to 3’. This procedure results in an ordered set of alignments necessary to detect ligation events between adjacent fragments. If paired end (PE) reads are analyzed, fragments from read1 and read2 are merged if their alignment intervals on the reference are within a specified distance in the correct order and if the same is true for all fragments that follow towards 3’. The resulting set of merged alignments is then classified into being of single or chimeric origin and if classified as a chimera, further checked to be self-chimeric or multi- chimeric according to the criteria listed in Table 2. Reads which do not fall in any of the listed classes, e.g. if only one of two PE reads can be mapped, are discarded.

**Table 2.** Classification criteria for reads.

| class | criterion |
| --- | --- |
| single | single alignment (same location for PE) |
| chimeric | split alignment with at least 2 parts |
| self-chimeric | split alignment with parts mapping to same annotation |
| multi-chimeric | split alignment with at least 3 parts |

### Ligation points

Each chimeric read is further analyzed and checked for ligation events. A ligation event is defined as a split alignment of two adjacent chimeric fragments on the same read with a user-specified maximal distance of a few nucleotides. The coordinates of the nucleotide closest to the partner fragment are defined as the ligation points of those fragments (Fig. 1b). Ligation points always sit on the edge of an aligned fragment on the side pointing towards the ligated partner. To identify ligation points, first, the read length chosen in the experiment must be sufficiently high to cover at least two fragments and second, the parameters for mapping the reads to the genome will define a lower boundary for the detection of fragments on the reads.

### Computation of complementarity

ChimericFragments computes the complementarity of two ligated fragments as a local alignment using the SW algorithm with a substitution matrix. Instead of representing mutation probabilities with the scores, we designed a matrix that roughly represents binding affinity between two nucleotides. The parameterization of the matrix can be chosen by the user. Predictions are made for two intervals of specified length around each ligation point of a ligated pair of fragments. To evaluate each base-pairing prediction, we assign a score s to it. For this, we take the complementarity score s_c_ from the SW algorithm and subtract it by a weighted shift score s_s_:

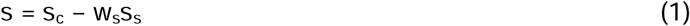

The shift score s_s_ is defined as the absolute difference of the ends of the complementarity regions found in the two sequences with respect to the observed intervals. This score can be used to select for base-pairing predictions which end at similar positions with respect to ligation points. The weight w_s_ can be defined by the user.

### Statistical evaluation

ChimericFragments implements two statistical evaluations of chimeras. First, we construct a null model by sampling complementarity scores s_i_ between random patches on the genome of specified length. The empirical cumulative density function ECDF of those scores s is computed and a p-value can be attributed to predictions:

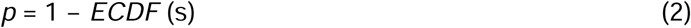

In general, multiple ligation events are found for a given pair of annotations. To summarize the events, we assess the distribution of the p-values for each pair and combine them by first computing the FDR with the method of Benjamini and Hochberg [70]. ChimericFragments offers multiple methods to combine the p-values from ligation points of the same interaction. Either by taking the minimum of the FDR values of all sampled ligation points for one interaction as a combined p-value to represent the probability of error when considering the strongest complementarity found, or by using the methods proposed by Fisher [71] or Stouffer [72], which both test against the null hypothesis of uniform distribution of p-values in the null model they are derived from. The FDR reported in the results tables is based on all combined p-values and is used to select interactions shown in the graph. To assess the possibility of detecting RNA-RNA interactions merely because both partners are close to each other or the chaperone protein of interest by chance, each pair gets assigned a statistical significance using Fisher’s exact test according to [26]. This significance is based on the number of times a pair was detected and the number of times, each partner was found in other interactions or as a single read.

### Visualization of RNA-RNA networks

ChimericFragments visualizes the network of interactions as a graph. The nodes in the graph correspond to annotations on the genome, and an edge represents all chimeras associated with the two annotations connected by the edge (Fig. 1d). Edges are directed and the direction of the edge represents the order, in which the fragments are found on their reads. An edge pointing from annotation A to annotation B indicates the fragment mapping to A was found further upstream on the read(s) than the fragment mapping to B. In our graphical representation of the network, the size of the nodes and edges correspond to the share of reads within the selected set of interactions. ChimericFragments is implemented as a web application based on Dash and Dash Bio [73] to support fast drawing and smooth repositioning of hundreds of nodes.

### Node positioning in graph drawings

To draw the RNA-RNA interaction network do without any prior positional information on the nodes of the graph, we use a technique called stress majorization [74] and chose the parameterization of this algorithm according to [75]. Stress majorization is applied to each connected component in the graph and the components are then placed with a simple guillotine bin packing [76] algorithm that fills a rectangle with all connected components while minimizing empty space between them. Together, those two techniques lead to reproducible graph drawings, which are similar for graphs with similar sets of nodes and edges.

### Plotting ligation points

Upon clicking on an edge in the graph, an interactive scatter plot of the ligation points in the coordinate frame of the annotations (Fig. 1e, Fig. S1b) is displayed. In this frame, +1 refers to the first nucleotide in the annotation, counting from -1 down in the upstream direction and from +1 up in the downstream direction. For merged annotations containing a CDS, +1 corresponds to the first nucleotide in the CDS. Each dot in the scatter plot shows the corresponding region of complementarity when hovered upon and size and color of the dot correlate with the number of supporting reads and the FDR associated with the prediction. An FDR-cutoff can be set which applies to FDR values computed from the p-values of the ligation points of currently selected interactions.

### Plotting aggregated complementarity

For every node in the graph, all predictions between the node and all of its partners with an FDR below a specified cutoff in the graph are aggregated in a summary plot (Fig. 1f, Fig. S1c). To compute this summary, for every position in the annotation all ligation points with overlapping regions of complementarity from all interactions in the current selection are summed up. Hovering upon the plotted data shows for every position in the corresponding annotation all partners with a complementary site at this position.

### Bacterial strains and growth conditions

All strains used in this study are listed in Supplementary Table S3. *V. cholerae* and *E. coli* strains were grown aerobically in LB medium at 37 °C. Where appropriate, antibiotics were used at following concentrations: 20 µg/mL chloramphenicol and 50 µg/mL kanamycin.

### Plasmid construction

All plasmids and DNA oligonucleotide sequences used in this study are listed in Supplementary Tables S2 and S4, respectively. GFP fusions were cloned as described previously[35] using previously determined transcriptional start sites [45, 46]. Inserts were amplified from *V. cholerae* genomic DNA with the respective oligonucleotide combinations indicated and cloned into linearized pXG10 vectors (KPO-1702/-1703) via Gibson assembly [77]; pSM002 (KPO-5251/-5252), pSM003 (KPO-5247/-5248), pJR004 (KPO-2460/-2462), pJG020 (KPO-8756/-8757) and pAL069 (KPO-9358/- 9359). For pNP040 (KPO-1832/-1833) and pNP045 (KPO-1838/-1839), pXG10 and respective inserts were digested with NsiI and NheI and ligated. The constitutive sRNA expression plasmid pAL062 (KPO-9233/-9234) was constructed by PCR amplification of the respective sRNA from *V. cholerae* genomic DNA and cloned into linearized pEVS143 vector [78] (KPO-0092/-1397) via Gibson assembly. Site directed mutagenesis of pJR006 using KPO-9373 and -9374 resulted in pAL077.

### Fluorescence measurements

To validate interactions captured by RIL-seq, GFP fluorescence measurements were performed as described previously [79] with *E. coli* Top10 cells cultivated overnight in LB medium. Cells were washed and resuspended in PBS and relative fluorescence was measured with a Spark 10 M plate reader (Tecan). Control strains not expressing fluorescent proteins were used to subtract background fluorescence.

### RNA isolation and Northern blot analysis

Total RNA sample preparation and Northern blot analyses were performed as previously described [80]. Membranes were hybridized in Roti-Quick buffer (Carl Roth) with [32P]-labelled DNA oligonucleotides at 42 °C. Signals were visualized using a Typhoon Phosphorimager (GE Healthcare). Oligonucleotides for Northern blot analyses are listed in Supplementary Table S4.

### Western blot analysis

Total protein sample preparation and Western blot analyses of 3XFLAG-tagged fusions were performed as previously described [81]. 3XFLAG-tagged fusions were detected using mouse anti-FLAG antibody (Sigma; F1804). RNAPα served as loading control and was detected using rabbit anti-RNAPα antibody (BioLegend; WP003). Signals were visualized using a Fusion FX EDGE imager and quantified with BIO-1D software (Vilber Lourmat).

## Supporting information

combined Supplemental Material

## DECLARATIONS

### Ethics approval and consent to participate

Not applicable.

### Consent for publication

Not applicable.

### Availability of data and materials

All datasets analyzed in this study are published and available online. Sequencing data of RNA interactome studies are available under the following accession codes: RIL-seq *E. coli* (ArrayExpress, E-MTAB-3910), RIL-seq *V. cholerae* (GEO, GSE198671), RIL-seq EPEC (ArrayExpress, E-MTAB-8806), RIL-seq *S. enterica* (GEO, GSE163336), RIL-seq *P. aeruginosa* (GEO, GSE216135), CLASH *E. coli* (GEO, GSE123050), LIGR-seq *B. subtilis* (ArrayExpress, E-MTAB-8490) and CLASH *S. cerevisiae* (GEO, GSE114680). Term-seq and dRNA-seq sequencing data can be found under the GEO accession codes “GSE144478“ and “GSE62084”, respectively. The source data used in this article is available at https://github.com/maltesie/ChimericFragmentsFigures (DOI: 10.6084/m9.figshare.24806094). The source code of ChimericFragments is available at: 10.6084/m9.figshare.24806097.

### Authors’ contributions

M.S. and K.P. designed the study; A.L. performed the experiments; M.S., A.L. and K.P. analyzed data; M.S. and K.P. wrote the manuscript.

## Acknowledgements

We thank Andreas Starick and Yvonne Greiser for excellent technical support. We thank Björn Voss (University of Stuttgart) for comments on the manuscript and all members of the Papenfort lab for insightful discussions and suggestions.

