## Supplementary material for "ChimericFragments: Computation, analysis, and visualization of global RNA networks": combined Supplemental Material

**This supplement contains:**

Supplementary Figure Legends

Supplemental References

Figures S1-9

Tables S1-4

**TABLE OF CONTENTS**

**Figure S1** Control elements of ChimericFragments

**Figure S2** Graph and table views of the graphical interface of ChimericFragments

**Figure S3** Additional plots view of the graphical interface of ChimericFragments

**Figure S4** Summary view of the graphical interface of ChimericFragments

**Figure S5** Benchmarking of bwa-mem2

**Figure S6** Results in additional datasets

**Figure S7** Basepairing characteristics

**Figure S8** Network of interactions between IGRs and mRNAs

**Figure S9** Network of interactions between IGRs and mRNAs

**Table S1** Validated basepairing sites in literature

**Table S2** Plasmids used in this study

**Table S3** Strains used in this study

**Table S4** Oligonucleotides used in this study

#### Supplementary Figures

##### Figure S1

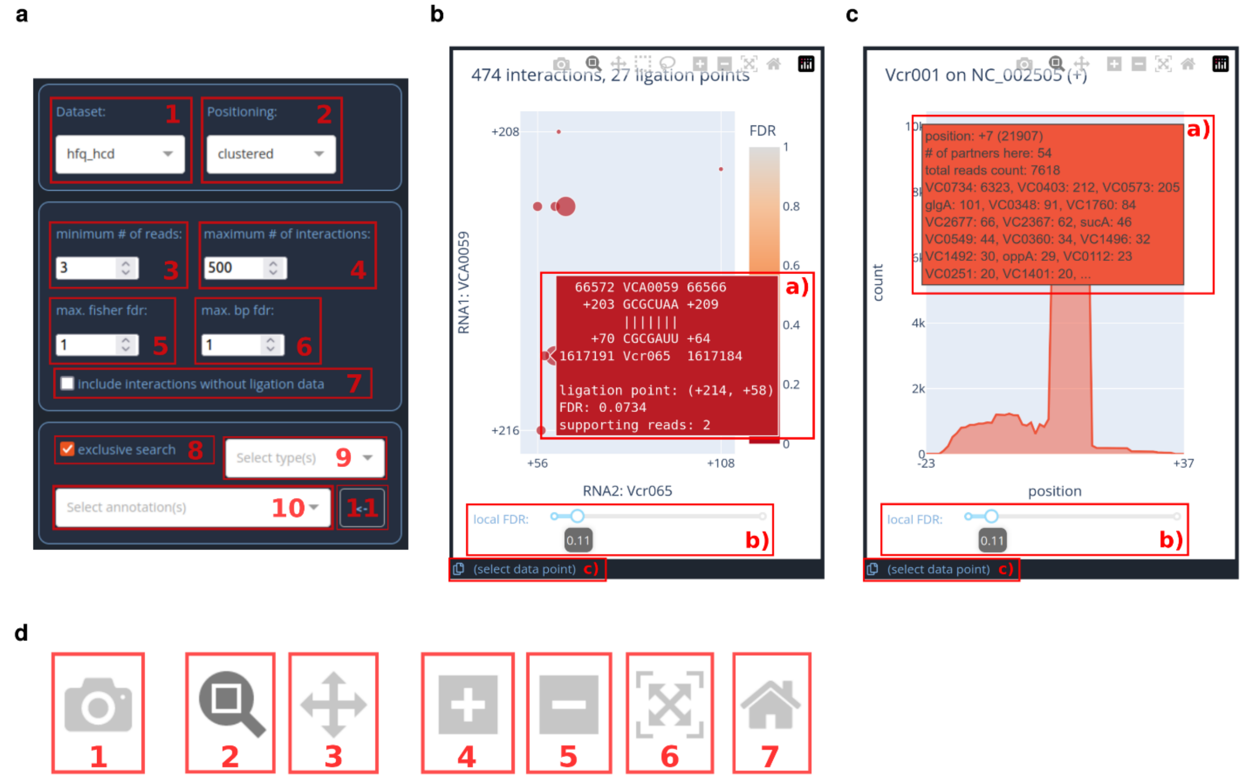


**Figure S1: Control elements of ChimericFragments**

**a,** Controls panel of the interface with multiple components for filtering and presenting the results of the computational part: 1) Choose between all analyzed conditions. 2) Choose a node positioning algorithm. “clustered” positions nodes based on stress majorization of the adjacency matrix of the connected components and packs the components with a guillotine bin packing algorithm. “grid” positions the nodes on an approximately squared grid sorted by the number of reads, each node was found in. 3) Choose the minimum number of reads per interaction. 4) Choose the maximum number of interactions in the network. 5) Only display interactions with a significance according to the Fisher exact test with the selected FDR-cutoff. 6) Only display interactions with a significance according to our complementarity test with the selected FDR-cutoff. 7) Choose, if interactions without ligation points are displayed. 8) select, if multiple searching criteria get combined exclusively, i.e. when two types are selected, exclusive search would only display interactions between those two, while non-exclusive search would select for interaction with at least one of the selected types involved. 9) select, which types of annotation to include in the selection. 10) select, which genes to include in the selection. 11) click to add a selected node in the network to the list of genes to display. **b,** Plot of ligation points for a selected interaction in the network. Dots represent ligation points at the coordinates in the respective annotations. RNA1 is on the y-axis, RNA2 on the x-axis. Dots are colored according to the FDR of the ligation point and sizes correspond to the number of reads this ligation point was found in. 1) Hovering over a dot will display a tooltip with a basepairing prediction and corresponding information. 2) The ligation points displayed in the plot can be filtered according to the local FDR (see Methods). 3) Clicking on a dot in the plot enables the clipboard tool. When enabled, clicking on the clipboard symbol will copy the content of the tooltip to the clipboard. **c,** Aggregation plots for all basepairing predictions of all interactions of a selected node. Only basepairing predictions with a local FDR lower than the global cutoff are aggregated. Predictions for interactions where the selected node was found as RNA1 in a pair (blue) are separated interactions where the node was found as RNA2 (red). 1) Hovering over any position in the plotted lines displays a tooltip with information on all targets predicted to bind at the respective position and corresponding information. 2) The ligation points displayed in the plot can be filtered according to the local FDR (see Methods). 3) Clicking on a dot in the plot enables the clipboard tool. When enabled, clicking on the clipboard symbol will copy the content of the tooltip to the clipboard. **d,** Menu for all plots in the graphical interface. 1) Save a png (portable network graphics) of the currently displayed part of the plot. 2) Switch to zoom mode. The plot will zoom into rectangular selections match the plot. 3) Switch to pan mode. 4) Zoom into the plot. 5) Zoom out of the plot. 6) Display all plotted content. 7) Return to initial view of the plot.

#### Figure S2

**a**

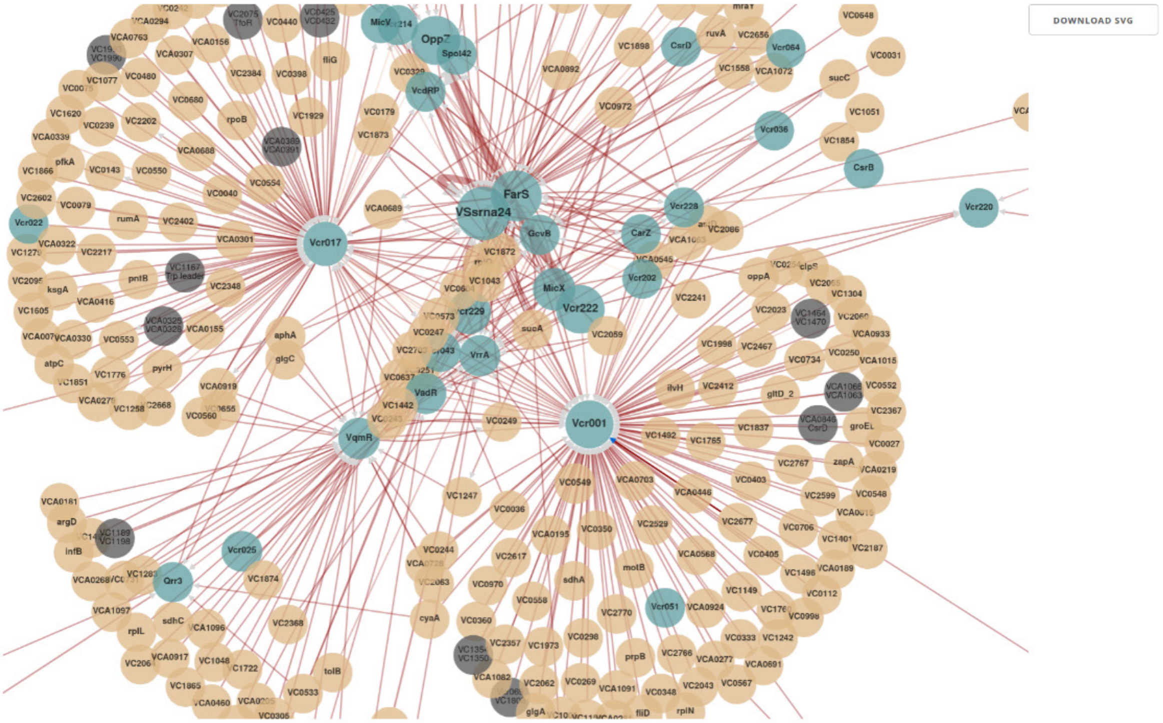


**b**

DOWNLOAD CSV

| name1 | type1 | name2 | type2 | nb_ints | fisher_fdr | odds_ratio | bp_fdr | in_libs |
|---|---|---|---|---|---|---|---|---|
| Vcr222 | sRNA | OppZ | sRNA | 101032 | 0 | 579.83 | 5.7148e-176 | 2 |
| FarS | sRNA | VSsrna24 | sRNA | 81190 | 1 | 0.016069 | 0.000088911 | 2 |
| Vcr229 | sRNA | VSsrna24 | sRNA | 22502 | 1 | 0.16835 | 2.6396e-32 | 2 |
| VSsrna24 | sRNA | FarS | sRNA | 16420 | 1 | 0.0033522 | 4.8368e-55 | 2 |
| VC0734 | CDS_UTRS | Vcr001 | sRNA | 16414 | 0 | 17.005 | 1.6299e-80 | 2 |
| Vcr025 | sRNA | VadR | sRNA | 12828 | 0 | 1286.5 | 8.8081e-7 | 2 |
| MicX | sRNA | VC0972 | CDS_UTRS | 12724 | 0 | 19.595 | 3.1673e-22 | 2 |
| VSsrna24 | sRNA | Vcr229 | sRNA | 10680 | 1 | 0.084127 | 7.9513e-124 | 2 |
| VC0549 | CDS_UTRS | Vcr001 | sRNA | 9526 | 0 | 350.65 | 2.34e-19 | 2 |
| VC0972 | CDS_UTRS | MicX | sRNA | 9459 | 0 | 14.708 | 7.8565e-43 | 2 |
| VC0573 | CDS_UTRS | Vcr001 | sRNA | 6730 | 0 | 136.46 | 1.3599e-55 | 2 |
| VCA0059 | CDS_UTRS | VrrA | sRNA | 5807 | 0 | 539.76 | 6.1636e-93 | 2 |
| oppA | CDS_UTRS | Vcr001 | sRNA | 4031 | 0 | 42.982 | 6.0603e-11 | 2 |
| VC1337 | CDS_UTRS | Spot42 | sRNA | 3978 | 0 | 68.364 | 0.0051365 | 2 |
| VC2086 | CDS_UTRS | Vcr001 | sRNA | 3936 | 0 | 19.845 | 3.6645e-8 | 2 |
| VSsrna24 | sRNA | Vcr001 | sRNA | 3684 | 1 | 0.023559 | 1.4601e-7 | 2 |
| VCA1097 | CDS_UTRS | VqmR | sRNA | 3321 | 0 | 80.55 | 1.0772e-9 | 2 |
| VC1461 | CDS_UTRS | Vcr096 | sRNA | 3083 | 0 | 25443 | 4.6704e-17 | 2 |
| VC1149 | CDS_UTRS | Vcr001 | sRNA | 3056 | 0 | 678.74 | 1.4902e-47 | 2 |
| FarS | sRNA | Vcr001 | sRNA | 3055 | 1 | 0.029483 | 0.0049328 | 2 |
| QrrX | sRNA | Qrr3 | sRNA | 2891 | 0 | 398.34 | 0.070359 | 2 |
| VC1872 | CDS_UTRS | FarS | sRNA | 2759 | 1 | 0.50615 | 4.0488e-60 | 2 |
| VC1898 | CDS_UTRS | VSsrna24 | sRNA | 2632 | 3.0747e-31 | 1.3853 | 1.0528999999999998e-83 | 2 |
| Vcr017 | sRNA | VSsrna24 | sRNA | 2288 | 1 | 0.019171 | 0.11 | 2 |
| rpoB | CDS_UTRS | Vcr017 | sRNA | 2212 | 0 | 151.8 | 1.5684e-62 | 2 |
| VC1043 | CDS_UTRS | FarS | sRNA | 2195 | 1 | 0.52255 | 1.11e-96 | 2 |
| Vcr229 | sRNA | Vcr228 | sRNA | 2057 | 0 | 3.8918 | 1.2459e-14 | 2 |
| VC2469 | CDS_UTRS | Vcr017 | sRNA | 1996 | 0 | 63.332 | 1.0744e-15 | 2 |
| VC0305 | CDS_UTRS | VqmR | sRNA | 1774 | 0 | 29.291 | 6.4768e-14 | 2 |

**Figure S2: Graph and table views of the graphical interface of ChimericFragments**
**a,** Screenshot of the network view of the graphical interface of ChimericFragments. Nodes are colored according to their annotation type (sRNA in cyan, mRNA in brown and IGRs in dark grey). Interactions are directed according to the order on the sequencing read the pair was found in, i.e. the upstream fragment points to the downstream fragment. Clicking on nodes and edges in the graph displays additional plots to the left of the network area. Click "DOWNLOAD SVG" button to download the displayed area of the graph as a svg-file (scalable vector graphics). Nodes in the graph can be rearranged by clicking and dragging them. **b,** Screenshot of the table view of the graphical interface of ChimericFragments. All interactions displayed in the network view are displayed here with detailed information. Click "DOWNLOAD CSV" to download a csv-file (comma separated values) of all displayed interactions.

#### Figure S3

a

b

c

d

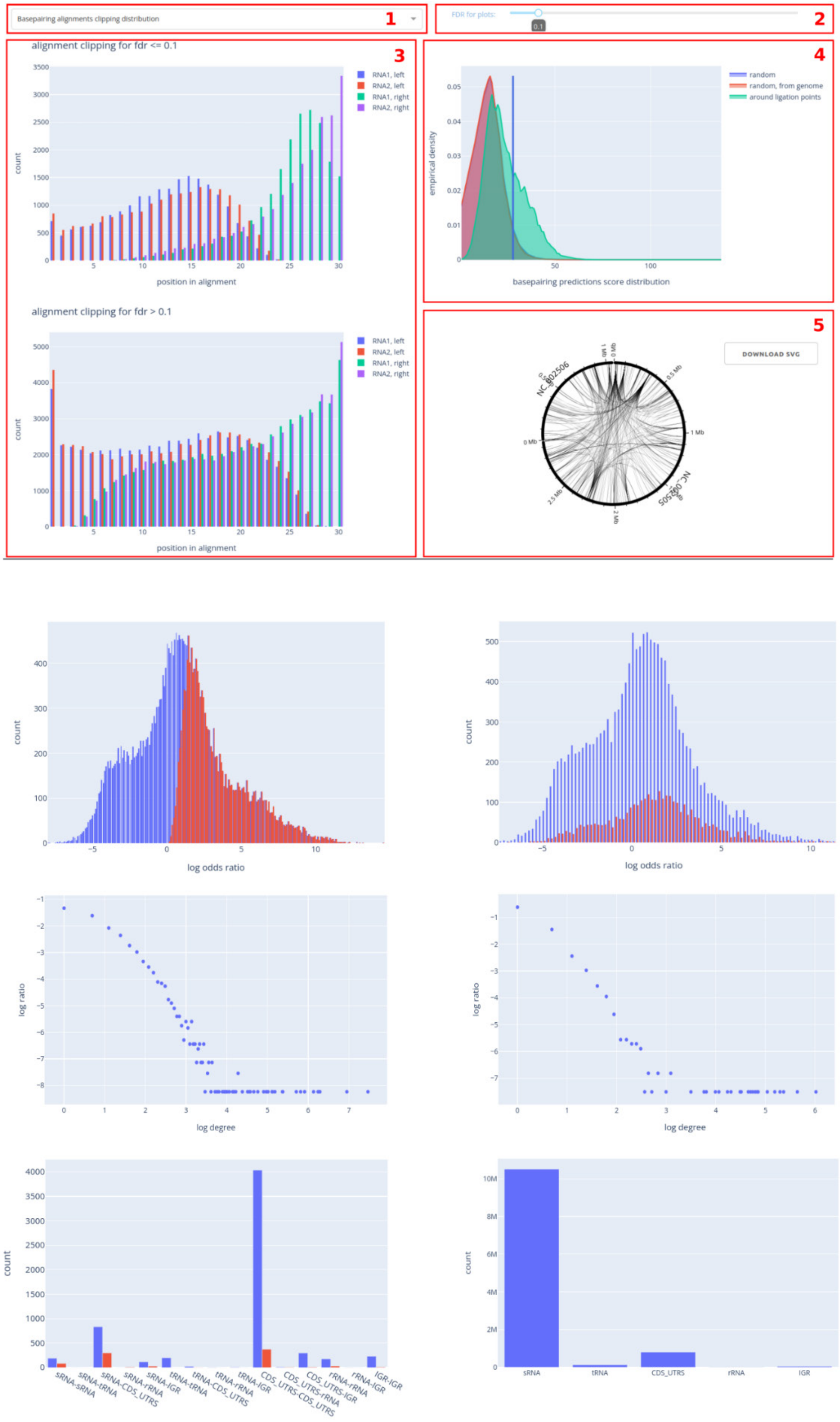

**Figure S3: Additional plots view of the graphical interface of ChimericFragments**

**a,** Additional plots view of the graphical interface of ChimericFragments. Four types of plots can be selected and interactively adjusted here. 1) Select between “Basepairing alignments clipping distribution”, “Odds ratio distribution”, “Annotation stats” and “Node degree distribution”. 2) Select FDR-cutoff for plots displayed in areas 3 and 4. 3) Area for plots selected in a. Here, the distribution of the end positions of the complementarity regions around ligation points is selected. For more detail, see main text and Figure 3. 4) Plot of the density of complementarity scores in a random model with equal probabilities for each base (blue), the random model used as null model in our statistical evaluation with random stretches of fixed length taken from the supplied genome (red) and the sequences around ligation points found in the selected dataset (green). The blue line corresponds to the selected FDR in b. 5) Circos plot of all interactions selected in the control panel and displayed in the network and table views (Extended Data Fig. 2 a and b). Click “DOWNLOAD SVG” to download the plot as a svg-file. **b,** Left: Odds ratio distribution of all interactions in the selected dataset. Significant interactions according to the Fisher exact test (red) with FDR cutoff selected with the FDR slider (panel a 2). Right: All interactions with a ligation point (blue) and significant interactions according to our complementarity test (red) with FDR cutoff selected with the FDR slider. **c,** Node degree distributions for networks filtered according to selected FDR. Left: Fisher exact test, right: our complementarity test. **d,** Bar plots of annotation types of interacting pairs.

**Figure S4**

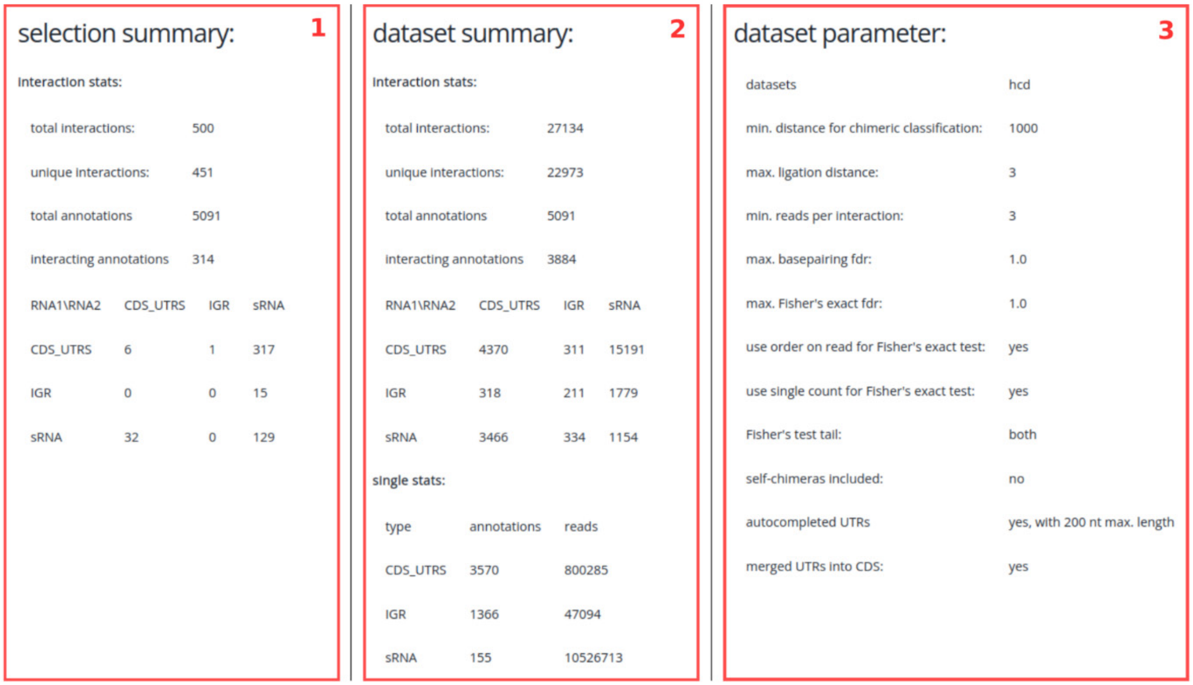


**Figure S4: Summary view of the graphical interface of ChimericFragments**
Screenshot of the summary view of the graphical interface of ChimericFragments. 1) Summary of the selected data displayed in the network and table views. 2) Summary statistics of the selected dataset. 3) Overview over some important parameters used in the analysis.

#### Figure S5

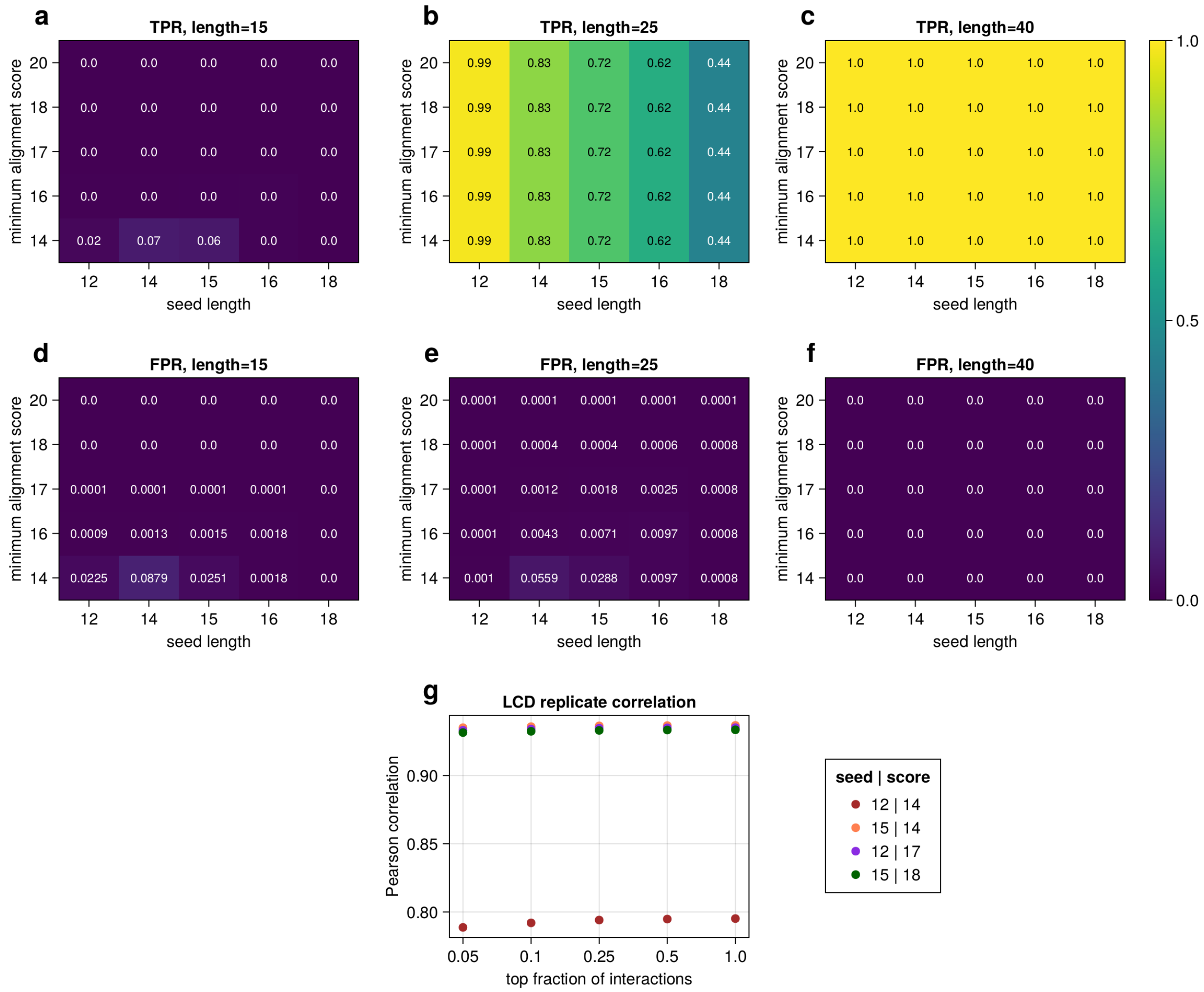


#### Figure S5: Benchmarking of bwa-mem2

**a-f,** true positive rate and false positive rate of alignments of chimeric reads of specified length in synthetic datasets with one mismatch (Table 1). For fragments of length 15, a maximum of 6% of the reads can be assigned correctly (**a**), while up to 11% get aligned to false positions in the genome (**d**). For fragments of length 25, the TPR ranges from 44% to 95% depending on the seed length (**b**) with FPRs from 0% to 6% (**e**). Fragments of length 40 get aligned with a TPR of 99% (**c**) and no false positives (**f**) independent of the choice of parameters. **g,** Pearson correlation of reads per interaction between the two replicates in the RIL-seq dataset.

#### Figure S6

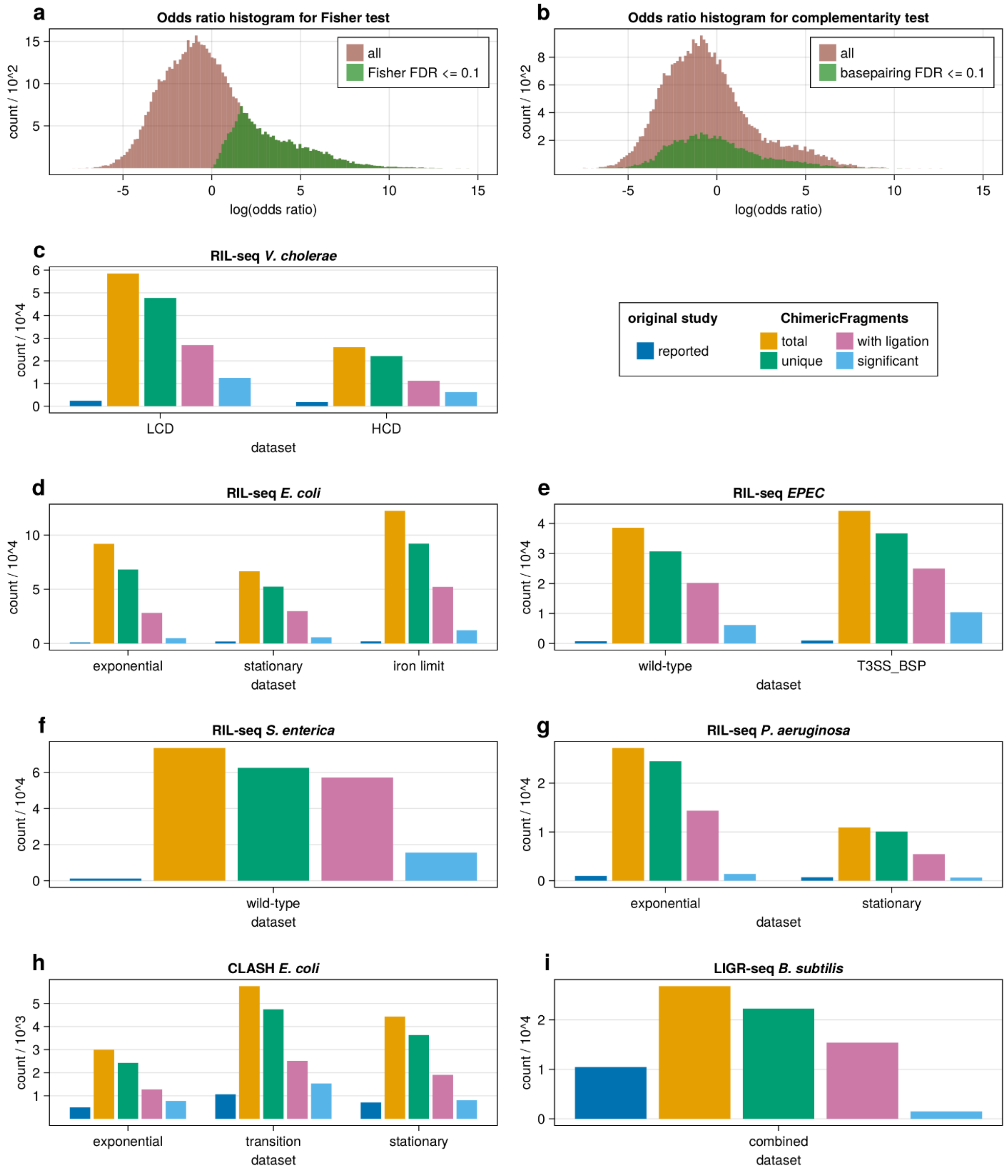


#### Figure S6: Results in additional datasets

**a,** Histogram of odds ratios for all interactions in the RIL-seq dataset (red) with significant odds ratios according to the right-tailed Fisher exact test with an FDR-cutoff of 0.1 (green). **b,** Histogram of odds ratios for all interactions in the RIL-seq dataset (red) with significant odds ratios corresponding to interactions which are significant according to our complementarity test with an FDR-cutoff of 0.1 (green). **c-i,** Number of interactions found in multiple published datasets with ChimericFragments. Shown are the reported number of interactions in the publication of the dataset (dark blue), the total number of interactions found by ChimericFragments (orange), the number of unique interactions found by ChimericFragments (green), the number of unique interactions with at least one ligation point (pink) and the number of unique interactions with a significant combined FDR (≤ 0.25, light blue, p-values combined with Stouffer's method). (**c**) RIL-seq experiment in *V. cholerae* (1) with a read cut-off of 3. (**d**) RIL-seq experiment in *E. coli* (2) with a read cut-off of 3. (**e**) RIL-seq experiment in *EPEC* (3) with a read cut-off of 3. (**f**) RIL-seq experiment in *S. enterica* (4) with a read cut-off of 3. (**g**) RIL-seq experiment in *P. aeruginosa* (5) with a read cut-off of 3. (**h**) CLASH experiment in *E. coli* (6) with a read cut-off of 1. (**i**) LIGR-seq experiment in *B. subtilis* (7) with a read cut-off of 3.

#### Figure S7

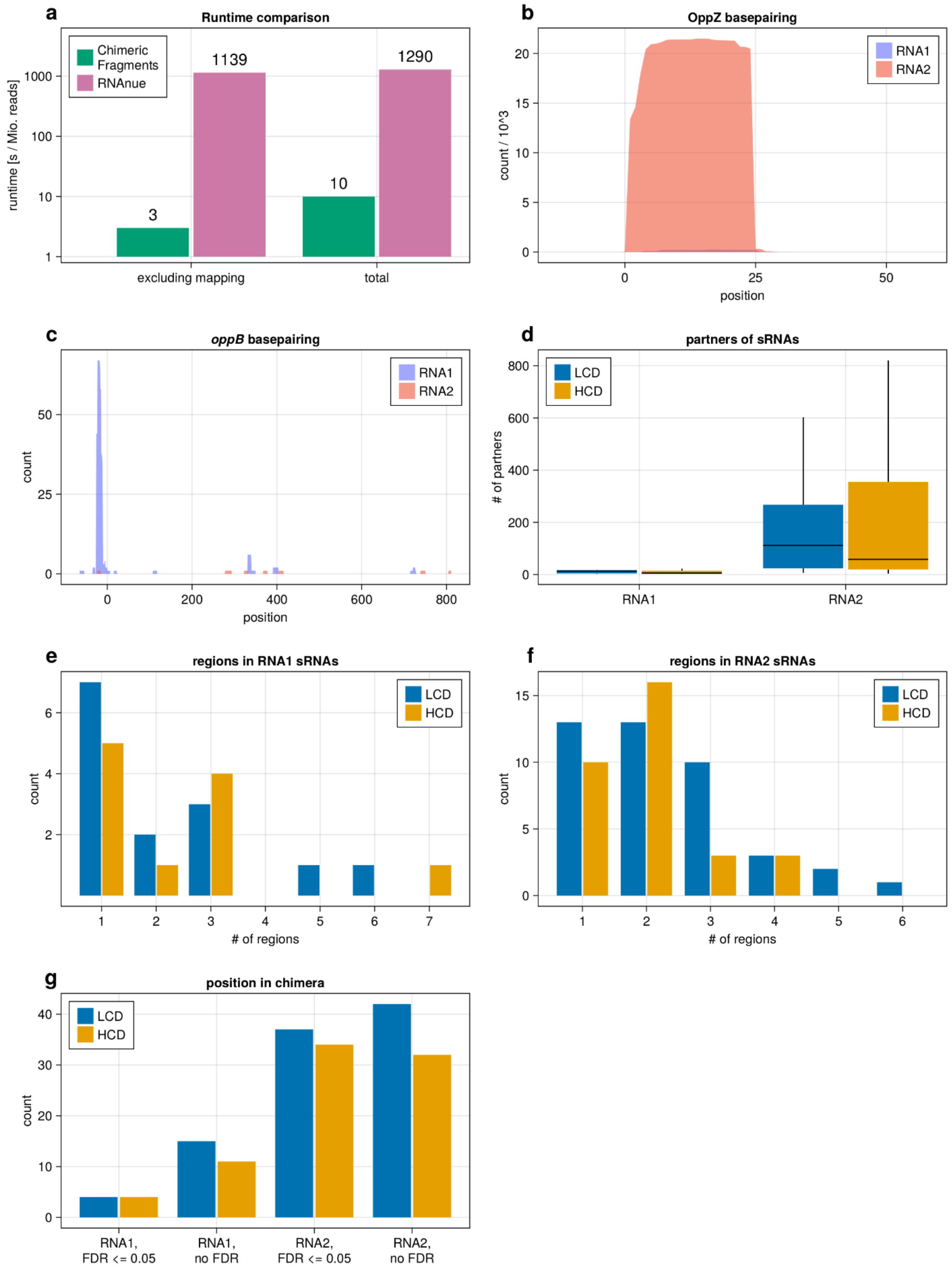

a
Runtime comparison
Chimeric Fragments
RNAnue
runtime [s / Mio. reads]
1000
100
10
1
3
1139
10
1290
excluding mapping
total
b
OppZ basepairing
RNA1
RNA2
count / 10^3
20
15
10
5
0
0
25
50
position
c
oppB basepairing
RNA1
RNA2
count
50
25
0
0
200
400
600
800
position
d
partners of sRNAs
LCD
HCD
### of partners
800
600
400
200
0
RNA1
RNA2
e
regions in RNA1 sRNAs
LCD
HCD
count
6
4
2
1
2
3
4
5
6
7
### of regions
f
regions in RNA2 sRNAs
LCD
HCD
count
15
10
5
0
1
2
3
4
5
6
### of regions
g
position in chimera
LCD
HCD
count
40
30
20
10
0
RNA1, FDR <= 0.05
RNA1, no FDR
RNA2, FDR <= 0.05
RNA2, no FDR

**Figure S7: Basepairing characteristics**

**a,** Runtime comparison between ChimericFragments and RNAnue. Both tools were run with default parameters for PE sequencing data of the RIL-seq experiment in *V. cholerae*(1). **b,** Aggregation of basepairing predictions with FDR < 0.25 for the OppZ sRNA. **c,** Aggregation of basepairing predictions with FDR < 0.25 for the *oppB* mRNA. **d,** Boxplot of number of partners per sRNA grouped by their class defined in (**g**) in two RIL-seq datasets obtained from low cell density (LCD) and high cell density (HCD) in *V. cholerae*. Bars correspond to 2 quartiles around the median, lines stretch to 1.5 times the distance from median to the quartile values. Outliers are omitted. **e-f,** Number of distinct regions found in the aggregated basepairing plots of each sRNA according to their class defined in (**g**). To compute the number of regions, the aggregation plots were normalized to sum to 1 across all positions and then iteratively positions with highest normalized counts were collected until certain cut-off was reached. The resulting set of positions was then queried for continuous intervals and they were counted. This procedure was repeated for cut-offs between 0.3 and 0.7 and the maximum resulting count was selected to represent the number of distinct regions. **g**, Number of sRNAs found in the RIL-seq experiment in *V. cholerae* with respect to the order they were found in with more targets (RNA1 for first in pair, RNA2 for second in pair), once for the whole dataset and once for all significant interactions (Fisher exact FDR ≤ 0.05).

**Figure S8**

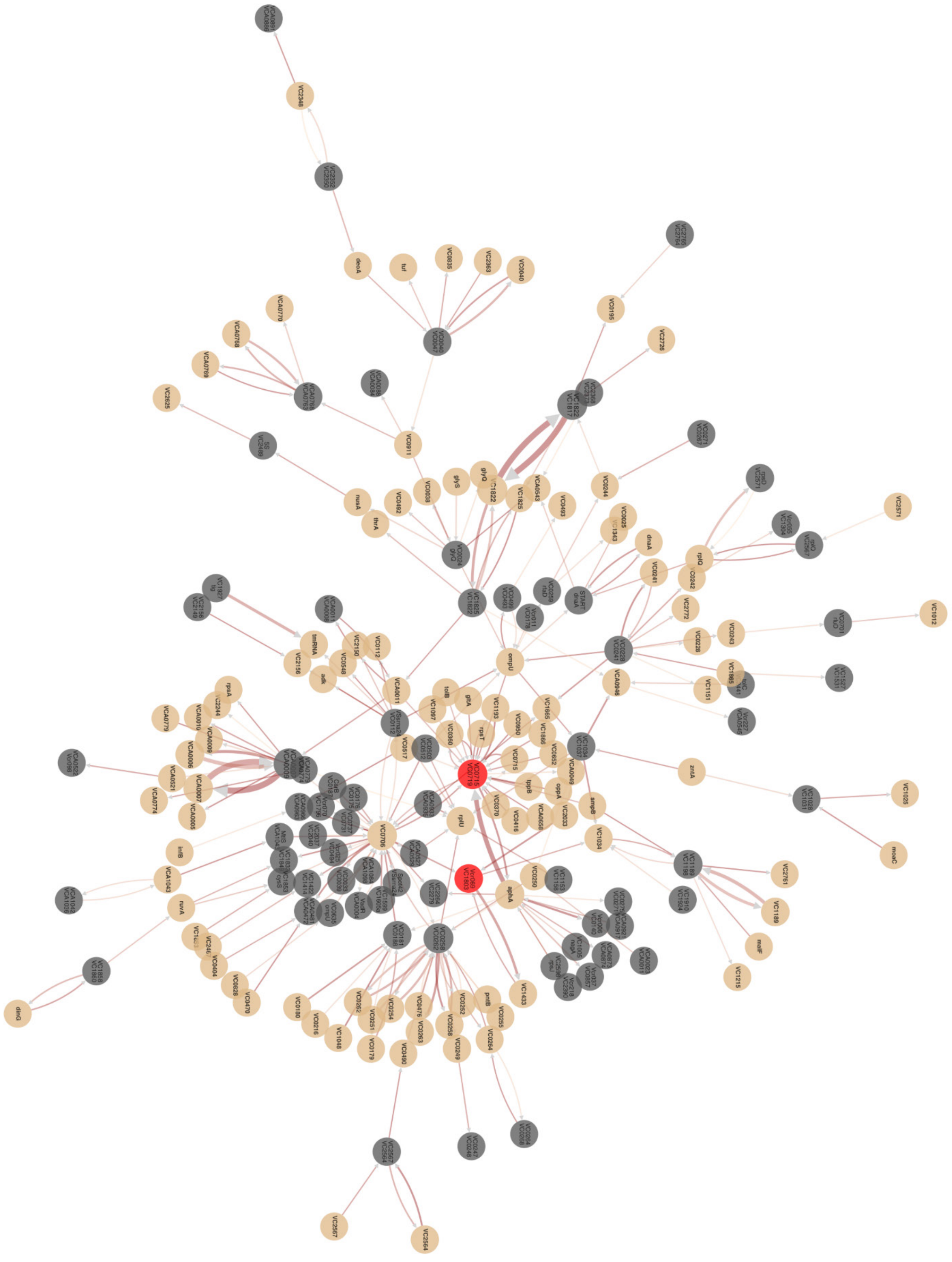

**Figure S8: Network of interactions between IGRs and mRNAs**

Largest connected component in the Graph of interactions between intergenic regions (IGRs, dark grey) and mRNAs (brown). IGRs VC0715:VC0719 and Vcr069:VC1803 are highlighted in red.

### Figure S9

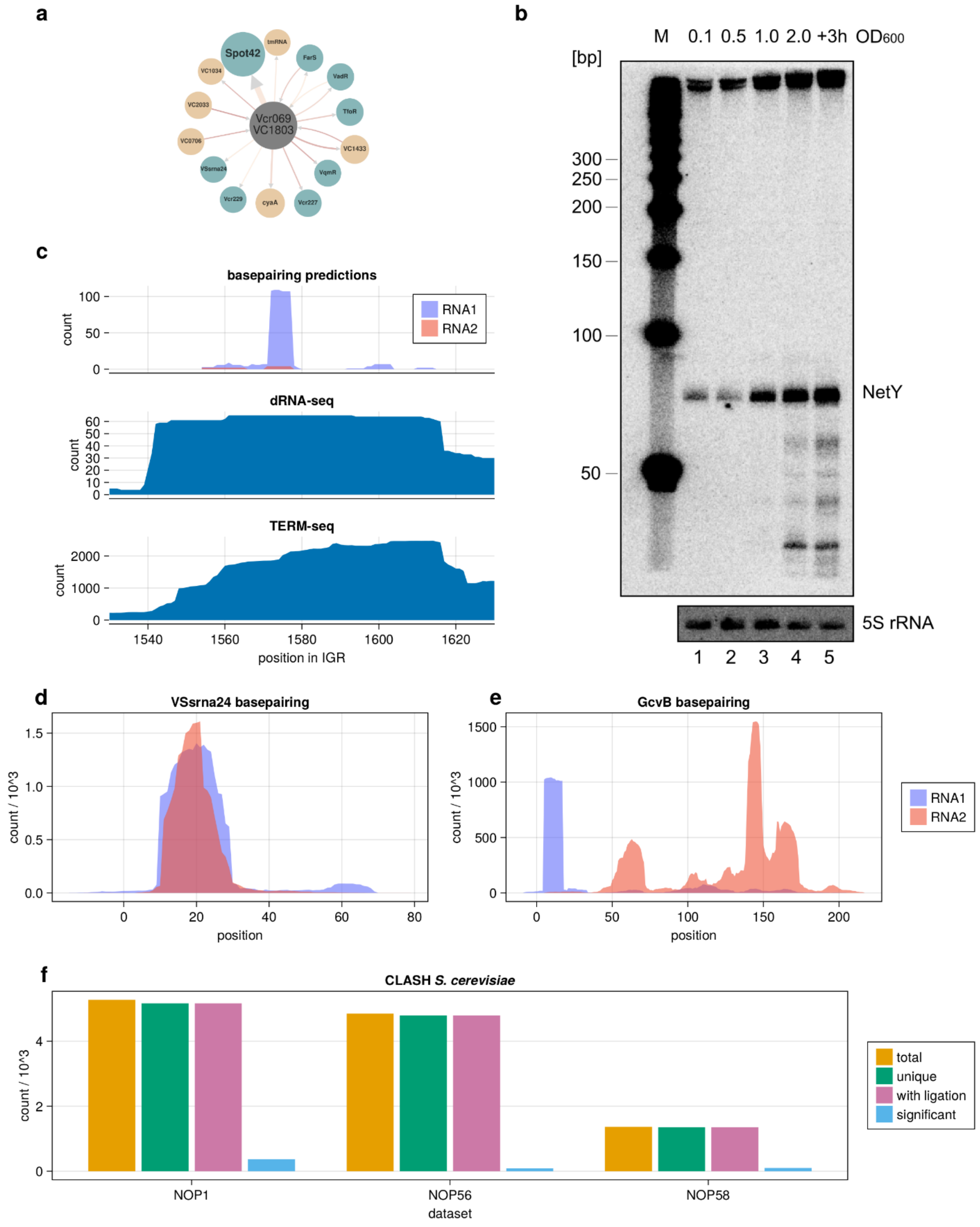
a
Spot42
tmRNA
FarS
VadR
TfoR
VC1433
VqmR
Vcr227
cyaA
Vcr229
VSsrna24
VC0706
VC2033
VC1034
Vcr069
VC1803
b
M 0.1 0.5 1.0 2.0 +3h OD600
[bp]
300
250
200
150
100
50
NetY
5S rRNA
1 2 3 4 5
c
basepairing predictions
RNA1
RNA2
count
dRNA-seq
TERM-seq
position in IGR
d
VSsrna24 basepairing
count / 10^3
position
e
GcvB basepairing
f
CLASH S. cerevisiae
total
unique
with ligation
significant
NOP1
NOP56
NOP58
dataset

**Figure S9: NetY and additional results**

**a,** Network of all interactions captured by RIL-seq of NetY sRNA. **b,** Aggregation of basepairing predictions with FDR < 0.25 for the IGR Vcr069:vc1803 sRNA in the RIL-seq dataset. Nodes are colored according to their annotation type (sRNA in cyan, mRNA in brown and IGRs in dark grey). **c,** NetY levels were monitored bv Northern blotting. RNA samples from *V. cholerae* wild-type cells were collected at various stages of growth. 5S ribosomal RNA served as a loading control. **d,** Aggregation of basepairing predictions with FDR < 0.25 for the VSsrna24 sRNA. **e,** Aggregation of basepairing predictions with FDR < 0.25 for the GcvB mRNA. **f**, Number of interactions found in a CLASH experiment in *S. cerevisiae* (8) with ChimericFragments. Shown are the total number of interactions found by ChimericFragments (orange), the number of unique interactions found by ChimericFragments (green), the number of unique interactions with at least one ligation point (pink) and the number of unique interactions with a significant combined FDR (≤ 0.25, light blue, p-values combined with Stouffer's method).

**Table S1: Validated regulatory interactions and basepairing sites in literature adopted from previous studies (9-11).**

Table S1 is available as an excel sheet.

#### Table S2: Plasmids used in this study

| Plasmid trivial name | Plasmid Stock name | Relevant fragment | Comment | Origin, marker | Reference |
|---|---|---|---|---|---|
| pXG10-sfGFP | pXG10-sfGFP | lacZ'::sfGFP | Template plasmid for translational reporter | pSC101*, $Cm^R$ | (12) |
| pXG10-*vc2085* (*sucC*) | pSM002 | 5'UTR + 20 aa of *vc2085* | Translational GFP reporter | pSC101*, $Cm^R$ | This study |
| pXG10-*vc0108* (*polA*) | pSM003 | 5'UTR + 20 aa of *vc0108* | Translational GFP reporter | pSC101*, $Cm^R$ | This study |
| pXG10-*vca0946* | pNP040 | 5'UTR + 15 aa of *vca0946* | Translational GFP reporter | pSC101*, $Cm^R$ | This study |
| pXG10-*vc2361* | pNP045 | 5'UTR + 15 aa of *vc2361* | Translational GFP reporter | pSC101*, $Cm^R$ | This study |
| pXG10-*vc1043* | pJR004 | 5'UTR + 20 aa of *vc1043* | Translational GFP reporter | pSC101*, $Cm^R$ | This study |
| pXG10-*vc0122* (*cyaA*) | pJR026 | 5' UTR + 20 aa of *vc0122* | Translational GFP reporter | pSC101*, $Cm^R$ | (9) |
| pXG10-*vca0166* | pJR036 | 5' UTR + 20 aa of *vca0166* | Translational GFP reporter | pSC101*, $Cm^R$ | (9) |
| pXG10-*vc0966* | pMH062 | 5' UTR + 20 aa of *vc0966* | Translational GFP reporter | pSC101*, $Cm^R$ | (9) |
| pXG10-*vc2030* (*rne*) | pJR045 | 5' UTR + 20 aa of *vc2030* | Translational GFP reporter | pSC101*, $Cm^R$ | (9) |
| pXG10-*vc1904* | pKT005 | 5' UTR + 20 aa of *vc1904* | Translational GFP reporter | pSC101*, $Cm^R$ | (9) |
| pXG10-*vc2602* | pJG020 | 5'UTR + 20 aa of *vc2602* | Translational GFP reporter | pSC101*, $Cm^R$ | This study |
| pXG10-*vc2647* (*aphA*) | pKP462 | 5' UTR + 20 aa of *vc2647* | Translational GFP reporter | pSC101*, $Cm^R$ | (13) |
| pXG10-*vca0848* | pAL069 | 5' UTR + 20 aa of *vca0848* | Translational GFP reporter | pSC101*, $Cm^R$ | This study |
| pCMW-1 | pCMW-1 | | Control plasmid | p15A, $Kan^R$ | (14) |
| p-*spot 42* | pAS001 | *spot 42* | *spot 42* expression plasmid | p15A, $Kan^R$ | (15) |
| pEVS143 | pEVS143 | P*tac* promotor | Constitutive overexpression plasmid (template) | p15A, $Kan^R$ | (16) |
| p-*netX* | pAL062 | *netX* | *netX* expression plasmid | p15A, $Kan^R$ | This study |
| p-*farS* | pJR006 | *farS* | *farS* expression plasmid | p15A, $Kan^R$ | (17) |
| p-*farS* G54C* | pJR014 | *farS* G54C* | *farS* G54C* expression plasmid | p15A, $Kan^R$ | (17) |
| p-*farS* G34C* | pAL077 | *farS* G34C* | *farS* G34C* expression plasmid | p15A, $Kan^R$ | This study |

#### Table S3: Strains used in this study

| Strain | Relevant markers / Genotype | Reference / Source |
|---|---|---|
| ***V. cholerae*** | | |
| KPS-0014 | C6706 wild-type | (18) |
| KPVC-12901 | C6706 *aphA::3XFLAG* | (9) |
| ***E. coli*** | | |
| Top10 | *F- mcrA Δ(mrr-hsdRMS-mcrBC) φ80lacZΔM15 ΔlacX74 nupG recA1 araD139 Δ(ara-leu)7697 galE15 galK16 rpsL(StrR) endA1 λ-* | Invitrogen |
| S17λpir | *ΔlacU169 (ΦlacZΔM15), recA1, endA1, hsdR17, thi-1, gyrA96, relA1, λpir* | New England Biolabs |

#### Table S4: Oligonucleotides used in this study

| Name | Sequence 5' to 3' | Description |
|---|---|---|
| KPO-0092 | CCACACATTATACGAGCCGA | Plasmid construction (pEVS143) |
| KPO-1397 | GATCCGGTGATTGATTGAGC | Plasmid construction (pEVS143) |
| KPO-1702 | ATGCATGTGCTCAGTATCTCTATC | Plasmid construction (pXG10) |
| KPO-1703 | GCTAGCGGATCCGCTGG | Plasmid construction (pXG10) |
| KPO-5251 | GAGATACTGAGCACATGCATCTCAGATTTCTGAGTAATG | Plasmid construction (pSM002) |
| KPO-5252 | CCAGCGGATCCGCTAGCCGGTACAGGCAAACCGAATTC | Plasmid construction (pSM002) |
| KPO-5247 | GAGATACTGAGCACATGCATGTTATCCACAGAATGTAGGGTTG | Plasmid construction (pSM003) |
| KPO-5248 | CCAGCGGATCCGCTAGCCGATAAAGGTAAGAGGAGCC | Plasmid construction (pSM003) |
| KPO-1832 | gttttATGCATATCCCAGAGAGGATGGTAAGA | Plasmid construction (pNP040) |
| KPO-1833 | gttttGCTAGCATCGCCGTACGCTTTACATAC | Plasmid construction (pNP040) |
| KPO-1838 | gttttATGCATAGCTCAACAAGATCTGAAAATTAAC | Plasmid construction (pNP045) |
| KPO-1839 | gttttGCTAGCCCAGATAGAGTTCAGTAGGTC | Plasmid construction (pNP045) |
| KPO-2460 | GATAGAGATACTGAGCACATGCATAGGATCAGCCACCACTGAG | Plasmid construction (pJR004) |
| KPO-2462 | CCAGCGGATCCGCTAGCTGTGGCTAGCGTTACCGTC | Plasmid construction (pJR004) |
| KPO-8756 | GATAGAGATACTGAGCACATGCAT AACTAGCAATCAGACTTGGAAAG | Plasmid construction (pJG020) |
| KPO-8757 | CCAGCGGATCCGCTAGC CGCGATACCAAGGATGTAACC | Plasmid construction (pJG020) |
| KPO-9233 | TCGGCTCGTATAATGTGTGGTTCGGTTGGCATGATCAGCA | Plasmid construction (pAL062) |
| KPO-9234 | GCTCAATCAATCACCGGATCAAAAGAAAAAGACGCGCTGAAAG | Plasmid construction (pAL062) |
| KPO-9358 | GAGCCAGCGGATCCGCTAGCTCCAAGATGGTGAAATAAGGTC | Plasmid construction (pAL069) |
| KPO-9359 | TAGAGGTACCGGTTGTTAACGTCGGCTCAATGAATACTGAT | Plasmid construction (pAL069) |
| KPO-9373 | CAGATTAGTTTCGCTAATCGAACAcACGCAGACT | Plasmid construction (pAL077) |
| KPO-9374 | CATGGGACAGAGTCTGCGTgTGTTCGATTA | Plasmid construction (pAL077) |
| KPO-9229 | CGTCATCGATTTACTGCCACG | NetX oligoprobe |
| KPO-9231 | GGCAACCAATGACCAACCTG | NetY oligoprobe |
| KPO-0243 | TTCGTTTCACTTCTGAGTTCGG | 5S oligoprobe |
